# ReMind: A Retrospective Self-Report Paradigm for Studying Mind-Wandering Onset During Reading

**DOI:** 10.64898/2026.05.14.725227

**Authors:** Haorui Sun, Alaina Birney, Niharika Singh, Ardyn Olszko, Peijin Chen, Jin Ke, Monica D. Rosenberg, David C. Jangraw

## Abstract

Mind-wandering (MW) is a frequent and pervasive phenomenon, yet it is commonly assessed using self-reports or probe-based methods that offer limited temporal precision regarding its onset. In this study, we introduce a novel paradigm, ReMind, that estimates the onset and duration of MW episodes during natural reading by combining retrospective self-reports with eye-tracking. Participants indicated the words where they believed their mind started and stopped wandering, and these reports were aligned with gaze timestamps to estimate MW onset. Using data from 44 participants, we examined whether knowledge of MW onset improves the detection of MW from eye-tracking signals. To evaluate relevance for both self-report and thought-probe paradigms, we additionally simulated thought probes by randomly sampling time points during reading. Logistic regression classifiers trained on eye-tracking features extracted from time windows anchored to MW onset achieved AUROC scores of 0.659 and 0.621 under the self-report and simulated thought-probe paradigms, respectively, using leave-one-subject-out cross-validation. In both cases, onset-aligned windows outperformed classifiers trained using arbitrary MW windows. Sliding-window analyses further revealed systematic temporal changes around MW onset, with classification performance peaking at approximately 3 seconds after onset. Feature-level analyses showed reduced fixation rate and fixation dispersion, along with increased pupil size following MW onset. Together, these findings characterize the temporal progression from on-task reading to MW. Overall, ReMind provides a useful framework for studying the temporal dynamics of MW during naturalistic reading.

## Introduction

Mind-wandering (MW), often described as task-unrelated thought (Giambra, 1989a; Smallwood & Schooler, 2006, 2015), is a ubiquitous and consequential phenomenon. Previous studies have shown that our minds wander frequently, with MW reported in approximately 30-60% of sampled moments during daily life (Kane et al., 2007; Killingsworth & Gilbert, 2010; Seli et al., 2018). Although people report generating innovative solutions or ideas during mind-wandering episodes (Mooneyham & Schooler, 2013), MW is most often associated with an unhappy mood (Killingsworth & Gilbert, 2010) and impaired task performance (Barron et al., 2011; Cotton et al., 2023; Van Vugt & Broers, 2016).

During reading, mind-wandering hinders comprehension (Ebbert et al., 2024; Farley et al., 2013; Mooneyham & Schooler, 2013; Smallwood, Fishman, et al., 2007; Smallwood et al., 2008; Unsworth & McMillan, 2013; Wammes et al., 2016). In tasks requiring sustained attention, such as driving or piloting, mind-wandering can lead to accidents or even fatalities (Galera et al., 2012; Gil-Jardiné et al., 2017; Gouraud et al., 2017). Despite its prevalence and varied consequences, the cognitive processes leading to the onset of MW remain unclear (Smallwood, 2013; Smallwood & Schooler, 2015). One major challenge is that MW onset is not directly detectable with existing methods, which typically capture only the moment participants become aware of their mind-wandering or are externally prompted to report it (Chu et al., 2023; Greve & Was, 2022; Schooler, 2002; Smallwood & Schooler, 2006). In this study, we aim to address this gap by introducing our ReMind paradigm using it in a free-viewing reading task to investigate the onset and temporal dynamics of MW during reading.

Prior studies of MW have primarily employed self-report methods (Chu et al., 2023) and/or thought probes (Greve & Was, 2022; Weinstein, 2018) to detect MW episodes. Self-reports rely on participants’ subjective awareness: whenever individuals notice that their minds have wandered away from the task, they are instructed to report the occurrence of MW (Schooler, 2002). In contrast, thought probes are random prompts to participants during a task, requiring them to indicate whether their thoughts are on-task or mind-wandering at each prompt (Giambra, 1995; Smallwood & Schooler, 2006). Both self-report and probe-based methods have been widely adopted in experimental paradigms across domains such as reading (Broadway et al., 2015; Faber et al., 2018; Reichle et al., 2010), sustained attention (Seli et al., 2016, 2018; Zanesco et al., 2021), and learning (Conrad & Newman, 2021; Farley et al., 2013), and they remain the dominant tools for identifying the occurrence of MW during task performance.

However, both methods fail to capture when MW begins. Instead, they tend to mark the moment when MW ends, either because a participant detected and reported the episode or because the thought probe interrupted it. In the case of self-reports, participants indicate MW only after they become aware that their attention has drifted, known as meta-awareness (Schooler, 2004). Because this meta-awareness is distinct from MW itself (Seli et al., 2017; Smallwood, McSpadden, et al., 2007), using this moment as a reference point can blur the line between when MW starts and when it is noticed or resolved. Thought probes, on the other hand, can occur at any point during a MW episode. Since the timing of probes is random and MW episodes vary in duration, it is difficult to determine when the episode began. For researchers trying to identify markers of upcoming MW episodes or model how MW unfolds over time, this ambiguity in timing poses a major challenge.

To infer the onset of MW, some studies have made assumptions such as attributing MW to the entire preceding trial (D’Mello et al., 2020; Groot et al., 2022; Stawarczyk et al., 2020) or using arbitrary time windows (Faber et al., 2018; Reichle et al., 2010). Although these approaches are often used out of necessity in the absence of more precise alternatives, they introduce measurement error that weakens the sensitivity and specificity of analyses. For example, when trials are long or vary in duration, assuming that MW spans the whole trial can obscure transitions between mental states and exaggerate the time spent in MW. Similarly, fixed windows, such as defining MW as the last 5 or 10 seconds before a report, impose artificial boundaries that may not reflect participants’ actual experience. These imprecise definitions of MW periods introduce noise and potential confounds, which may help explain the contradictory findings across the literature (Kam et al., 2022; Steindorf & Rummel, 2020). This highlights the need for more accurate methods of estimating MW onset: precise timing is essential not only for understanding how MW unfolds moment-to-moment, but also for testing key hypotheses about the cognitive processes that underlie it.

To address this methodological gap, we developed a novel approach, ReMind, that combines retrospective self-report with eye-tracking during a free-viewing reading task. During the task, we ask participants to self-report their MW episodes, and, critically, to indicate the specific words where they believe each episode began and ended. Reading provides an ideal context for studying MW: it is ecologically valid and in-lab tasks closely resemble natural reading experiences, especially when participants are allowed to progress at their own pace without time constraints or enforced viewing patterns. Moreover, MW occurs frequently during reading and has been widely studied in this domain (Bonifacci et al., 2023; Schad et al., 2012; Schooler et al., 2004; Smallwood, 2011). Prior research has observed that participants often engage in re-reading after MW episodes identified via self-report or thought probes, suggesting that they can retrospectively locate where their attention began to drift (Varao-Sousa et al., 2017). This behavior implies that readers are often able to identify the point in the text where MW started and use it as an anchor for re-engagement. Because eye movements are continuously recorded, we can precisely match this reported MW onset word with fixation timestamps. ReMind thus provides a more precise and natural estimate of when MW onset occurs compared to previous paradigms while maintaining ecological validity.

Prior work on MW during reading has identified signatures of mind-wandering from eye-movements, pupil size, and blink rates (Grandchamp et al., 2014; Mills et al., 2021; Reichle et al., 2010; Steindorf & Rummel, 2020). However, these analyses also tend to rely on noisy estimates of MW onset and duration. To identify more precise gaze-related measures of MW episodes and dynamics, we continuously collected eye-tracking data during our paradigm. This not only enables precise alignment with reported MW onsets but also provides an independent, non-invasive, and high-resolution measure of attentional dynamics. Eye tracking data can thus serve as a test of convergent validity for self-reported MW onset. If participants can accurately report their MW onsets, we might expect to see a change in one or more of these eye-tracking metrics at the reported onset time.

To this end, we collected eye-tracking data from participants as they completed our task during reading. We hypothesized that ReMind would enable more accurate estimation of MW onset compared to existing methods, thereby improving MW detection. To evaluate the effectiveness of our approach, we trained machine learning classifiers to distinguish MW from on-task states using eight eye-tracking features previously associated with MW in the literature. We benchmarked performance against standard paradigms that assume the mind-wandering episode began 2 or 5 seconds before the self-report (or thought probe) or at the start of the page. Using a sliding-window analysis, we also characterized the temporal profile of gaze changes surrounding MW onset. Together, our findings reveal the fine-grained dynamics of how MW begins and unfolds during reading and demonstrate that ReMind provides a powerful new framework for precisely identifying MW onset and tracking its progression over time.

## Methods

We report how we determined our sample size, all data exclusions (if any), all manipulations, and all measures in the study (Simmons et al., 2012).

### Participants

We recruited 58 participants from a university in the northeastern United States who could read English fluently and had no family history of neurological disorders or epilepsy. All participants underwent screening and provided informed consent before participating in the study. The study protocol was approved by the university’s Institutional Review Board.

14 participants were excluded due to issues such as difficulty working with the recording equipment, not completing all experimental runs, having only monocular eye-tracking data, or missing demographic information. This left a final sample of 44 participants. Their ages ranged from 18 to 64 years (median = 20, mean = 22.6). The group included 30 female, 9 male, and 5 non-binary participants, with 37 identifying as right-handed and 7 as left-handed (see Supplementary Table 1).

Given the final sample size of 44 participants, a sensitivity analysis conducted in G*Power (Faul et al., 2009) indicated that, for a two-tailed Wilcoxon signed-rank test with 𝛼 = .05 and power (1 − 𝛽) = .80, the study was sensitive to effects larger than 𝑑 = 0.44.

### Reading Materials

We selected five articles retrieved from Wikipedia in 2015 that participants were unlikely to already know but required no prior knowledge for comprehension. The topics were Pluto (the dwarf planet), the Prisoner’s Dilemma, Serena Williams, the History of Film, and the Voynich Manuscript. To standardize the reading material, we removed images and unnecessary jargon from the original text and divided each story into 10 pages. Each story contained approximately 2,200 words, resulting in about 220 words per page. We used a custom Python script to render this text into reading images for the task. Each page consisted of precisely 16 lines of black text in Courier font, displayed on a gray background on a computer screen.

To motivate participants’ reading and assess their comprehension, we developed a multiple-choice question for each page. We attempted to design comprehension questions so that each one could be answered correctly only if the reader had paid attention on that specific page. For example, after reading the sentence, “…Special effects were introduced and film continuity, involving action moving from one sequence into another, began to be used…”, participants might be asked, What does film continuity mean?, with options such as: (A) The plot of the film was fluent; (B) The film was played continuously without any pause; (C) Action moving from one sequence into another; (D) Motion pictures were produced without sound; and (E) I am not sure. At the end of each article, participants answered all the multiple-choice questions in randomized order, then rated the understandability of each story (“I was able to read and understand the text and questions”) and reported their familiarity with its content (“I had prior knowledge that helped me answer the questions”) on a 5-point Likert scale.

### Retrospective Self-Report (ReMind) Paradigm

The reading task was programmed using PsychoPy (Peirce et al., 2019), a platform for developing psychological experiments. Each experimental session consisted of five runs, one for each article. Articles were presented in a randomized order. Before the first run, participants received task instructions, including an explicit definition of mind-wandering (MW):

> *Let’s define ‘mind-wandering’ as any thoughts that make it so you couldn’t remember something you read. Mind-wandering could be: Totally random (e.g., “What did I have for lunch?”); Related to the text (e.g., “I wonder where this town is?”); Very brief (e.g., “I’m tired.”); Blank (e.g., “”); Useful (e.g., “I should remember that.”)*

Mind-wandering during reading is a familiar experience for most people. We defined it as fundamentally grounded in task-unrelated thought (Giambra, 1989a; Smallwood & Schooler, 2006). However, we framed it within our reading-for-comprehension task, where participants’ goal was to understand and remember the text. Accordingly, task-unrelated thoughts were defined as thoughts that interfered with reading comprehension or memory encoding. We also provided specific examples to help participants understand the definition and apply it during the task.

Participants read at their own pace with no time constraints per page. They were not permitted to re-read previous pages to ensure that only-pass reading behavior was captured. If participants noticed themselves mind-wandering, they were instructed to immediately press a designated key (“F”) on the keyboard. Upon reporting mind-wandering, participants were redirected to a dedicated reporting screen, which is a key feature distinguishing our paradigm from traditional self-report designs. On this screen, they used the mouse to click on the words where they believed their mind-wandering episode began and ended. The selected region was highlighted to help participants better visualize their attentional lapse. If they believed the mind-wandering began on the previous page, they were instructed to click the first word on the current page to indicate this. After submitting the retrospective self-report, participants returned to the same page to resume reading. For simplicity, each page allowed only one mind-wandering report.

We adopted a self-report rather than thought-probe approach to preserve the natural flow of reading and avoid interrupting ongoing cognitive processing. Although probe-caught methods reduce missed reports, externally imposed probes may disrupt the natural unfolding of both reading and mind-wandering. Unlike discrete attention tasks such as breathing counting (Levinson et al., 2014) or image-flashing paradigms (Pelagatti et al., 2025), reading is a continuous behavior that unfolds over time. We therefore aimed to preserve the integrity of the reading process and capture mind-wandering as naturally as possible within an ongoing reading context. This approach more closely approximates real-world reading, in which attentional lapses are detected through meta-awareness and followed by re-engagement with the text.

An additional advantage of preserving complete mind-wandering episodes is that it allowed us to simulate a probe-caught approach by randomly selecting time points within the reported mind-wandering period to mimic externally imposed probes. This enabled us to examine how knowledge of mind-wandering onset influences both self-report and thought-probe style analyses.

Before the main task, participants completed a short practice session using three pages from a different story to familiarize themselves with the retrospective reporting procedure. If they did not report any MW during these practice pages, they were shown the self-report screen on the final page for practice purposes. In addition, they answered three comprehension questions related to the practice story to preview the post-task assessment format.

Following the practice, participants began the main task and read through the 10 pages of their first article. Upon completing the article, participants answered 10 multiple-choice comprehension questions and the understandability and familiarity questions described above. This procedure was repeated for a total of five runs, with short breaks provided between runs.

### Eye-Tracking Data Acquisition

The experiment was conducted in a soundproof room measuring 7 × 7 ft to minimize distractions. The task was displayed on a Dell (model U2417H) monitor. The screen was 53.5 × 30.7 cm (1920 × 1080 pixels) and was placed 71 cm from the participant, so that each letter subtended 0.30 ° × 0.27 ° visual degrees, and lines were 1.44 ° apart. This makes each letter roughly equal to the resolution of the eye tracker (typical accuracy 0.25°-0.5°). The mouse and keyboard were arranged according to the participant’s handedness, placed within comfortable reach. To minimize eye movements and reduce artifacts in later analyses, right-handed participants were instructed to place their left index finger on the F key (used to report mind-wandering) and their left thumb on the spacebar (used to continue to the next page) throughout the reading. Left-handed participants used their right hand for this. To help maintain consistent eye tracking, participants were asked not to move or look at their hands during the task, but they were permitted to look down at the keyboard when responding to comprehension questions using the number keys. We simultaneously recorded eye-tracking and EEG data while participants performed the task; however, only eye-tracking data are reported in this paper.

We used an EyeLink 1000 Plus system for eye tracking (SR Research, Ottowa, Canada). The camera was positioned in front of the monitor (camera to screen distance of 16 cm), and a chin rest was used to stabilize participants’ heads during recording. The distance between the monitor and the camera as well as the distance between the camera and the participant’s eyes were recorded and entered into the SR Research software for calibration and reference. Binocular eye movements and pupil area were recorded at a frequency of 1000 Hz.

Participants underwent eye-tracking calibration and validation every run. We repeated a 9-point calibration until validation results were marked by EyeLink as good (worst point error < 1.5°, avg error < 1.0°). Once that was complete, we started recording their eye movements and EEG signals while they performed the reading task. PsychoPy sent triggers to both eye-tracking and EEG software at the onset of each page to synchronize the streams of data.

### Eye-Tracking Data Preprocessing and Feature Calculation

Fixations, saccades, and blinks were automatically detected using EyeLink software with default thresholds (saccade_velocity_threshold = 30; saccade_acceleration_threshold = 8000; saccade_motion_threshold = 0.1) and were parsed into separate data frames. Fixations were then mapped to specific words on the screen using the spatial locations of both fixations and words.

Pupil size data immediately before and after blinks are often unreliable due to the occlusion of the pupil and associated signal loss during eyelid closure. To address this, we applied a linear interpolation method (Hershman et al., 2018) to estimate pupil size during blinks. Since the rapid downward and upward eye movements at the start and end of a blink are typically classified as saccades by the EyeLink parser, we identified the saccades associated with each blink and used the pupil size values at the onset and offset of those saccades to linearly interpolate across the blink period. This approach reduced blink-related artifacts and produced a smoother, more continuous pupil size signal throughout the reading task.

We extracted a broad set of eye-tracking features (see Table 1 for detailed description of each included feature). We selected these features because they have been previously reported as correlates of MW or internal thought processes (Benedek et al., 2017; D’Mello et al., 2020; Grandchamp et al., 2014; Huang et al., 2019; Reichle et al., 2010). While other features from the literature where also considered, to reduce redundancy and improve model interpretability, we excluded features that were mathematically redundant such as those that were linear transformations of each other. For example, during reading, fixation and saccade counts are tightly coupled because each fixation is typically followed by a saccade, and fixation duration is often inversely related to fixation count when measured within fixed-length windows. We also considered blink-related features, such as blink frequency and duration, which have been included in some MW classification models (Grandchamp et al., 2014; Smilek et al., 2010).

**Table 1.** Summary of Eye-Tracking Features Used in the Analysis.

| Feature | Description |
| --- | --- |
| Fixation Rate | Fixation count on words per second. |
| In-Word Regression | The proportion of fixations within the same word relative to all fixations. |
| Out-Of-Word Regression | The proportion of fixations on previously read words relative to all fixations. |
| Word Frequency-Fixation Correlation | Correlation between word frequency (Zipf) and first fixation duration. |
| Fixation Dispersion | Average distance of each fixation from fixation center. |
| Vergence | Difference in gaze positions (left vs. right) weighted by fixation duration. |
| Pupil Size | Mean pupil size normalized by individual baseline. |
| Pupil Change Rate | Rate of change in pupil size over the time window. |

However, blinks occur relatively infrequently during reading (Abusharha, 2017; Cornelis et al., 2025). In our task, some participants exhibited long inter-blink intervals, resulting in missing values for short analysis windows and complicating model training. Therefore, we excluded blink-related features from our analyses.

### Analytical Framework

The primary goal of this analysis was to identify eye-tracking features that differentiate MW episodes from engaged reading and to evaluate whether precise estimates of MW onset improve detection. Our approach consisted of four main stages. First, we identified MW episodes using participants’ retrospective self-reports and constructed matched control intervals from non-MW reading. This step allowed us to define the temporal boundaries for subsequent feature extraction. Next, we extracted eye-tracking features from these intervals using several time-window definitions. These included windows aligned with the reported MW onset and fixed-length windows commonly used in previous studies. We then evaluated whether these features could distinguish MW from on-task reading by training logistic regression classifiers using a leave-one-subject-out cross-validation scheme.

Classification performance was quantified using the area under the receiver operating characteristic curve (AUROC) and compared across window types using permutation-based statistical tests. Feature importance was estimated with permutation methods to identify which eye-tracking measures contributed most to classification accuracy. Finally, to examine the temporal dynamics of MW, we applied a sliding-window analysis aligned to MW onset and self-report. This approach enabled us to track classifier performance over time and assess how individual features evolved as participants transitioned into and out of MW. This framework provided a comprehensive evaluation of MW detection, tested the utility of retrospective MW onset estimates, and offered insight into the time course of gaze-based signatures of MW.

### Mind-Wandering Episode Identification

We identified mind-wandering episodes based on participants’ self-reported onset and offset words, which are the words they clicked to indicate when they believed their mind started and stopped wandering. To do this, we looked for the sequence of consecutive fixations that occurred between those two reported words. In most cases, participants’ gaze stayed within the highlighted MW region defined by these words, but sometimes their gaze moved in and out of the region. To handle these cases, we used Kadane’s algorithm (Bentley, 1984) to find the segment with the highest number of consecutive fixations inside the reported window (see Supplementary Figure 1). The idea is simple: each fixation gets a positive value if it’s inside the region and a negative value if it’s outside, and the algorithm finds the longest stretch of fixations by maximizing the total sum of these values. The timestamps corresponding to the first and last fixation in this sequence were then taken as the MW onset and offset, respectively. This approach allowed us to handle ambiguous cases automatically and consistently (see Supplementary Figure 2).

### Behavioral Characteristics of Mind-Wandering

We examined whether mind-wandering reports varied as a function of reading page, article order, and story topic. MW was coded as a binary outcome for each page, indicating whether the participant reported MW on that page. We fit a generalized linear mixed-effects model with a binomial distribution and logit link function.

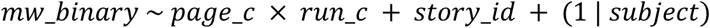

Fixed effects included centered page number, centered run order, story identity, and the interaction between page number and run order. Page number was centered around the grand mean to test whether MW changed over time within each article. Run order was centered around the grand mean to test whether MW changed across the experimental session. The interaction between page number and run order was included to test whether MW varied as a function of overall task time across the experiment. Story identity was included as a categorical predictor, with one story serving as the reference level. A random intercept for participant was included to account for repeated observations from the same participant.

In addition to mind-wandering frequency, we characterized the temporal properties of self-reported MW episodes. Specifically, we analyzed (1) MW duration, defined as the time between the self-reported onset and offset of a MW episode; (2) latency to MW onset, defined as the time between page onset and the reported MW onset; and (3) reporting delay, defined as the time between the reported end of a MW episode and the actual button press used to indicate the report. Distributions of these measures were visualized and summarized descriptively across participants and pages.

### Feature Window Definition

To evaluate the utility of our MW onset estimates and compare them with approaches used in prior work, we extracted eye-tracking features using several types of time windows (Figure 2B). We first defined our self-reported MW window, spanning from the participant’s reported MW onset to the moment of self-report. For each MW trial (i.e., time window), we identified a paired control trial using a window of equal duration that begins at the onset of a randomly selected fixation on a reading page without a MW report. We also implemented time windows commonly used in previous studies examining MW (Faber et al., 2018; Reichle et al., 2010). Specifically, we included fixed-length windows of 2 and 5 seconds, defined as the final 2 seconds (“2S”) or 5 seconds (“5S”) before the MW report. For control trials, these time windows were aligned to the end of non-MW pages. In addition, we extracted features from a full-page window (“FP”), extending from the start of the reading page to the report moment or page end. Because MW-reported pages typically ended earlier than non-MW pages, resulting in longer time windows for non-MW reading, we duration-matched each non-MW time window to a MW time window to ensure comparability in the time intervals used for feature calculation.

**Figure 1.**
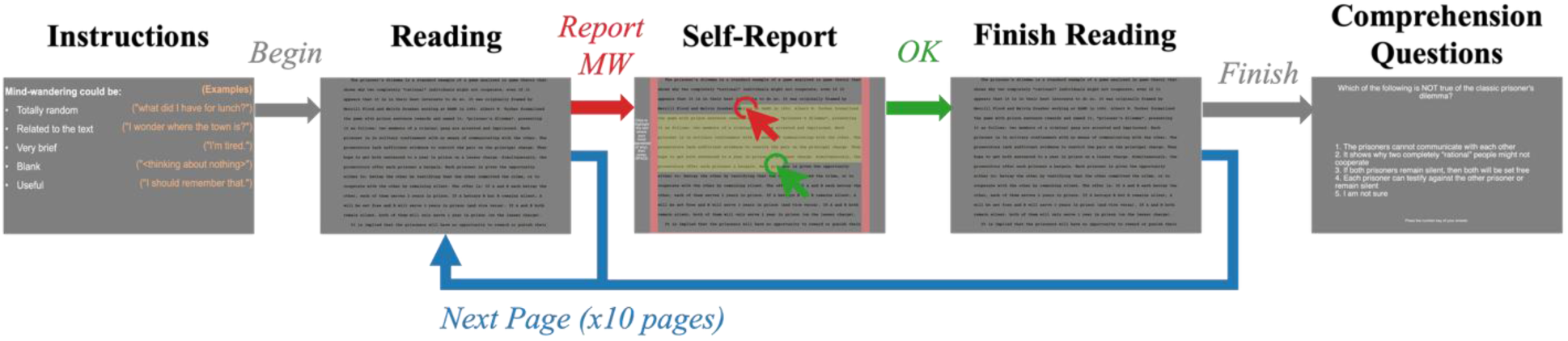
ReMind task paradigm. *Note.* Participants read 10 pages per run and reported mind-wandering by pressing a key, then clicking on the words where they believed their mind began and ended wandering. After reporting, they resumed reading from where they left off. Upon completing all 10 pages, participants answered 10 comprehension questions, one for each page.

**Figure 2.**
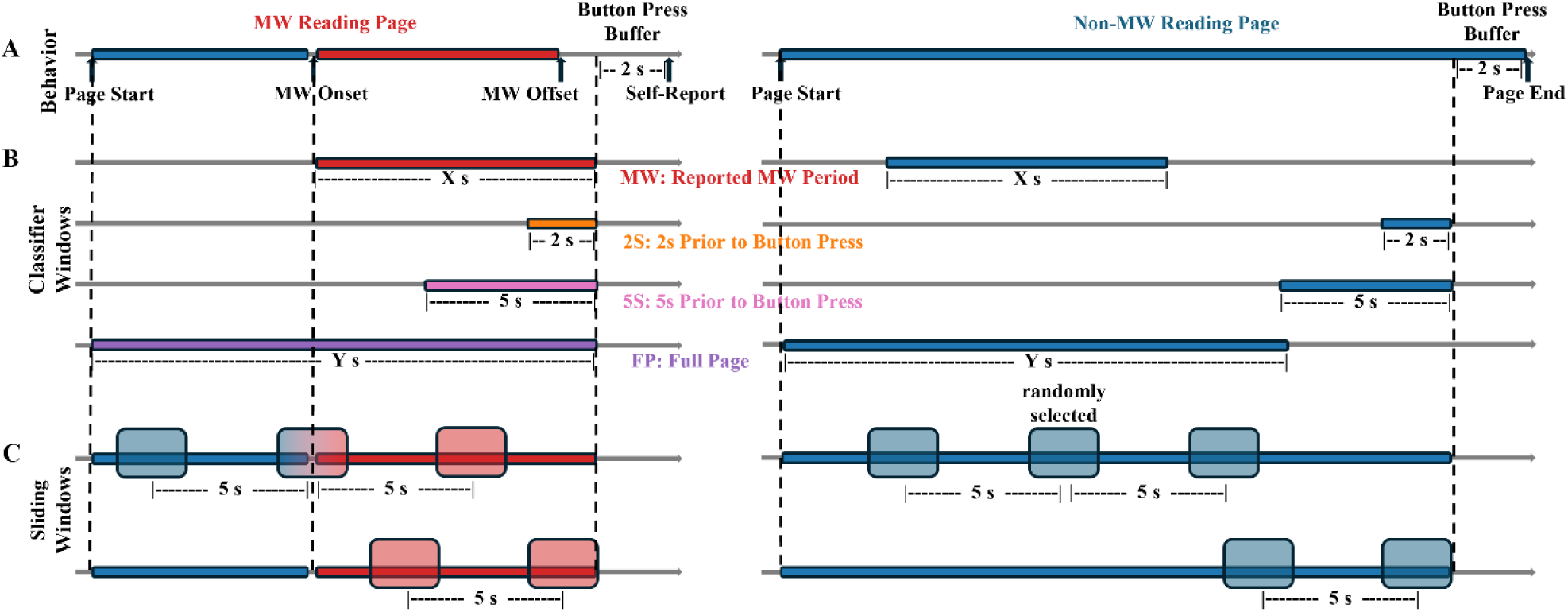
Time schemes for all analyses used in this study. *Note*. (A) Timeline of participants’ reading and reporting MW behavior. Each line represents one reading page. Red (left) indicates the reported mind-wandering (MW) period and blue (right) indicates normal reading. The last 2 seconds of each page were labeled as the button-press period and excluded from eye-tracking feature calculations. (B). Examples of the four classifier window types (MW, 2S, 5S, FP) during reading. Blue represents non-MW windows (control), while the other colors represent MW windows corresponding to each classifier type (colored to match the results seen in Figure 4). Non-MW and MW windows were matched in length. (C). Examples of sliding-window approaches for extracting eye-tracking features near MW onset and self-reports. Control sliding windows were applied in the same way to non-MW reported pages.

Because thought-probe paradigms are widely used in mind-wandering research, we also sought to examine how knowledge of MW onset would influence analyses within that framework. To do so, we simulated a thought-probe paradigm using our dataset (see Supplementary Figure 3). Specifically, for each reported MW episode, we randomly selected a time point within the duration of the episode and treated that moment as an externally imposed thought probe. The critical difference between the self-report and simulated thought-probe approaches was therefore the endpoint of the analysis window: self-report windows ended at the participant’s self-reported detection of MW, whereas simulated thought-probe windows ended at randomly sampled time points within the MW episode. Similarly, random probe moments were selected from non-MW reading periods for control trials. These sampled time points served as the endpoint of the corresponding analysis window, mimicking how thought probes interrupt ongoing reading and capture participants’ attentional state at a particular moment in time. For each simulated probe moment, we extracted features using the same set of window definitions (i.e., MW, 2S, 5S, FP) described above.

We observed that fixation-based estimates of MW offset did not consistently align with the moment participants pressed the button to report MW, frequently occurring earlier than the self-report (see Supplementary Figure 4). In addition, eye-tracking signals near the end of the page were less stable and exhibited abrupt changes (see Supplementary Figure 5), introducing potential confounds. Using fixation-based offsets would have introduced variability in the interval between MW offset and page end across trials, potentially distorting the temporal structure of our analyses. To maintain consistency with prior literature (Chu et al., 2023; Reichle et al., 2010) and ensure a stable reference point, we used the self-report button press as the endpoint of MW time window during eye-tracking feature calculation.

Furthermore, we excluded the final 2 seconds (Faber et al., 2018) of each page when defining time windows for feature extraction to minimize contamination from response-related activity and ensure that the analyzed intervals reflected ongoing cognitive processes rather than motor preparation or task-switching artifacts.

### Mind-Wandering Classification Analysis

We used a logistic regression classifier (implemented in Scikit-learn) to evaluate whether precise MW onset information improves the ability to distinguish between mind-wandering and on-task states.

Logistic regression was selected due to its simplicity, interpretability, and demonstrated effectiveness in capturing feature-based differences in attention (Brishtel et al., 2020a; Faber et al., 2018; Salazar et al., 2012). Before training, all eye-tracking features were standardized to have zero mean and unit variance. Standardization was performed separately for the training and testing sets to prevent data leakage and ensure valid evaluation.

To assess how well the model generalized across individuals, we used a leave-one-subject-out (LOSO) approach. Similar to a standard train-test split procedure, the model was trained and evaluated across multiple folds. However, in each fold, the model was trained on data from all participants except one and tested on the held-out participant. This process was repeated until every participant had served as the test subject exactly once.

LOSO evaluation is particularly important for this type of analysis. Because eye-tracking features were computed from temporal windows, LOSO helps prevent train-test leakage by ensuring that all testing samples came from separate recording sessions. It also provides a conservative and more accurate estimate of classifier generalizability across individuals. Specifically, LOSO reduces the influence of participant-specific differences in feature distributions that may remain even after normalization. Without participant-level separation, individual differences in reading behavior and mind-wandering tendencies could become confounding factors, allowing the model to partially rely on subject identity rather than eye-tracking patterns associated with mind-wandering.

We recorded the predicted probabilities and ground-truth labels for each test fold (i.e., participant) and computed the area under the receiver operating characteristic curve (AUROC) as the primary metric of classification performance. Instead of calculating AUROC separately for each fold, we aggregated predictions across all held-out test folds and computed a single overall AUROC score. This approach was chosen because some participants contributed relatively few samples, making participant-level AUROC estimates unstable and highly sensitive to small sample sizes. Averaging AUROC scores across folds could therefore artificially inflate or deflate the final performance estimate.

This classification pipeline was applied to datasets constructed using each of the window types described above (MW, 2S, 5S, and FP). Statistical comparisons between window types were performed using a permutation test with 200 iterations, in which mind-wandering labels were randomly shuffled to generate a null distribution of AUROC differences.

To better understand which eye-tracking features contributed most to classification performance, we examined feature importance using the permutation importance method provided in Scikit-learn. This approach measures the decrease in classifier performance when a given feature is randomly shuffled (repeated five times, as per the default setting of the permutation_importance function), while keeping the model and all other features unchanged. We repeated this procedure within each LOSO fold and averaged the results across participants to obtain group-level estimates of feature importance.

### Temporal Window Analysis

To examine how eye-tracking features evolve in relation to MW, we took advantage of our precisely timed onset and self-report data. For MW trials, we aligned the analysis to either the MW onset or the self-report moment. For control trials, onset-aligned windows were created by selecting a random fixation from a non-MW page and using its timestamp as a control onset. For offset-aligned analyses, the page end served as the control comparison point (Figure 2C).

We applied a sliding window approach to capture the time-resolved dynamics of each eye feature. For MW onset, we used a 2-second window stepped in 0.25-second increments across a 10-second interval centered on the onset (−5 to +5 s). For self-report, we analyzed only the 5 seconds preceding the report moment, excluding all post-report data due to the change in task demands (i.e., participants began clicking to identify MW onset and offset words). All sliding windows were constrained to fall entirely within the start and end of the reading page. Given the 2-second window length and the exclusion of the final 2 seconds of each trial, the last usable window ended at 2 seconds before the report, with its midpoint at 3 seconds before the report.

All eye-tracking features, except for pupil size, were calculated within each sliding window using standard aggregation (e.g., counts, averages) or correlation methods. Because pupil size is recorded continuously by the eye tracker, we treated it differently to preserve its native temporal resolution. Instead of averaging pupil size across the entire window, we extracted the instantaneous pupil size at the center time point of each window. This approach allowed us to maintain consistency in the number and alignment of trials across features, while preserving the fine-grained temporal structure of pupil dynamics.

### Time-Resolved Classification and Feature Dynamics

To evaluate how well MW could be detected at different time points, we trained a separate logistic regression classifier for each time point in the sliding window analysis. This allowed us to track classifier performance (AUROC) as a function of time relative to the MW onset or self-report. Since sample availability varied by time point due to differences in MW episode durations and window constraints (i.e., limitations imposed by how close the MW onset occurred to the beginning of the page), we identified the time point with the fewest valid samples and downsampled the other time points accordingly. We repeated this process separately for onset-aligned and report-aligned datasets. Classifier performance was tracked across time using AUROC scores at each time bin.

To further explore the transition of the eye feature set from MW onset to continuation, we trained a classifier at the 0-second mark (MW onset) and applied its weights across all other time points. This produced predicted probabilities (raw decision scores) that we visualized as a function of time. This approach enabled us to investigate whether MW onset and its continuation reflect the same underlying cognitive state (at least to the extent it can be observed through eye tracking features).

Finally, we visualized the temporal evolution of individual eye-tracking features around MW onset and self-report. For each time point, we used Wilcoxon signed-rank tests to compare feature values between MW and control conditions, using paired data for each participant. All p-values were corrected for multiple comparisons using the false discovery rate (FDR) procedure (Benjamini & Hochberg, 1995).

## Results

### Reading Comprehension and Prior Knowledge

Figure 3A shows an individual example of gaze traces and fixations during reading, with and without reported MW. Behavioral results suggested that the text was unfamiliar but understandable. Participants took a mean ± standard deviation of 58.98 ± 15.33 seconds to read each page (Figure 3B). On 181/220 (82.27%) of runs, participants agreed or strongly agreed that they could understand the text. On 17/220 (7.72%) of runs, participants agreed or strongly agreed that they had prior knowledge that helped them answer the comprehension questions. Participants achieved a mean comprehension accuracy of 61.82%, including 16.8% where they selected “I don’t know”. A one-sample t-test confirmed that this performance was significantly above chance levels of 25% across all story categories (*p* < .001).

**Figure 3.**
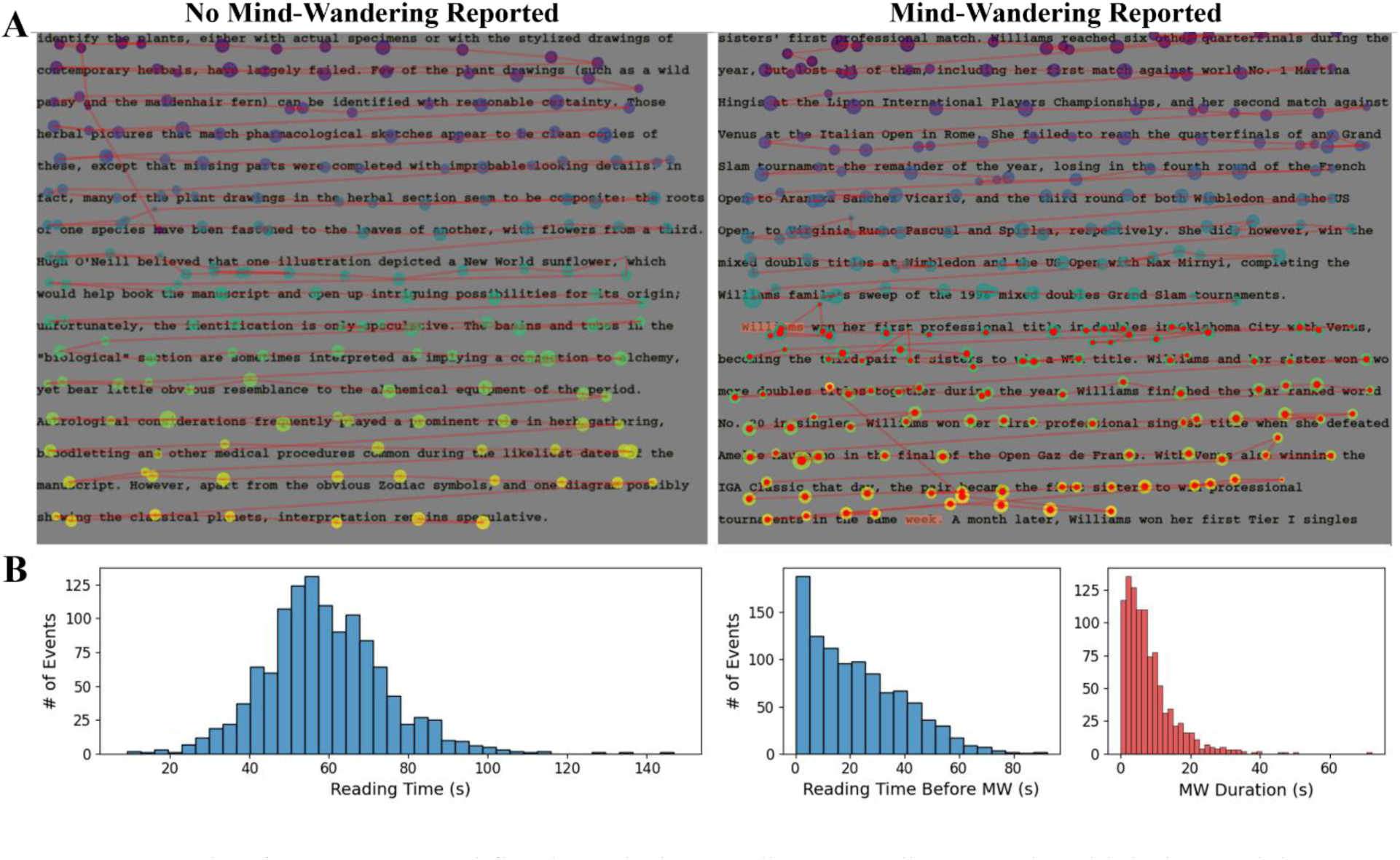
Reading behavior and task timing information. *Note*. (A) Example of gaze traces and fixations during reading: a reading page in which the participant read mindfully without reporting MW (left) and a reading page where the participant self-reported an MW episode (right). The onset and offset words of the MW episode are marked in red. Colored dots represent fixations. Larger dots indicate longer durations, and the color gradient from dark purple to yellow reflects the temporal order of fixations (from early to late) on the page. Fixations occurring within the self-reported MW episode are additionally center-colored in red. Consecutive fixations are connected by red lines, indicating saccades. (B) Histograms showing distributions of three timing measures: total reading time per page (left), reading time prior to MW onset (middle), and MW episode duration (right). MW episode duration here was defined as the reading time between the self-reported onset and offset times.

### Mind-Wandering Frequency and Duration

44 participants reported a total of 1001 mind-wandering episodes, which is 45.5% of all 2200 reading pages. The generalized linear mixed-effects model revealed a significant positive effect of page number, indicating that participants were more likely to report MW on later pages within an article, β = 0.037, SE = 0.017, z = 2.21, p = .027. There was no significant main effect of run order, β = −0.050, SE = 0.035, z = −1.44, p = .149, suggesting that MW rates did not reliably change across the five article runs. The interaction between page number and run order was also not significant, β = −0.005, SE = 0.012, z = −0.43, p = .665.

Story topic significantly influenced MW reports. Compared with the reference story, participants were significantly less likely to report MW during the Pluto article, β = −0.492, SE = 0.154, z = −3.20, p = .001. No significant differences were observed for Prisoners Dilemma, Serena Williams, or The Voynich Manuscript relative to the reference story.

The estimated standard deviation of the participant-level random intercept was 1.14, indicating substantial individual variability in overall MW tendency. The proportion of MW episodes per participant ranged from 0.04 to 0.90 (2 to 45 pages out of 50), with a standard deviation of 0.24 (Supplementary Figure 6).

Based on their reports, we computed the time interval between the beginning of a new page and the onset of MW. Participants began experiencing mind-wandering a median of 18.90 seconds after starting a new page. We also computed the duration of each MW episode as the time interval between the reported onset and offset. The median MW episode duration was 6.14 seconds, and the mean was 7.99 seconds. The distribution of these timing measures is shown in Figure 3B.

### Improved Classifier Performance Using Self-Reported MW Onset

To examine whether knowing MW onset improves classification, we trained logistic regression classifiers on eye-tracking features extracted from different time windows and compared classifier trained on features from MW windows aligned to the self-reported onset with those trained on more arbitrary time windows commonly used in prior work. Classifier performance is shown in Figure 4A. The classifier trained on features from self-reported MW episodes achieved the highest performance (AUROC = 0.659, random chance = 0.5), significantly outperforming the full-page (FP) condition (AUROC = 0.577, *p* < .01). Classifiers trained on the last 2- and 5-second windows prior to self-report achieved AUROC scores of 0.628 and 0.653, respectively. While slightly lower than the MW-onset window AUROC, these differences were not statistically significant (*p* = 0.125 and 0.465, respectively).

**Figure 4.**
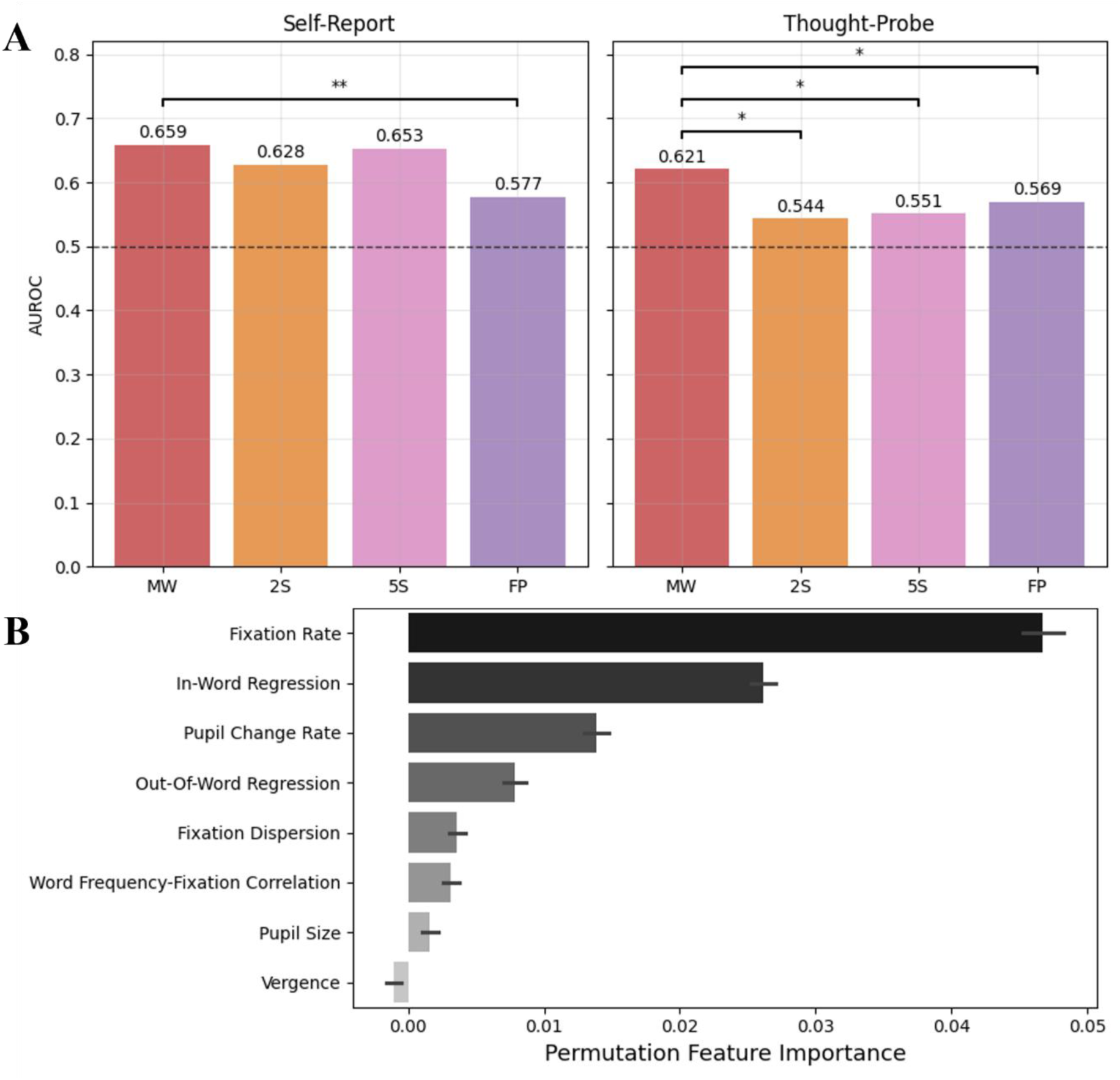
Logistic regression classifier performance and feature importance. *Note*. (A) AUROC scores for classifiers trained using different time windows under the self-report and simulated thought-probe paradigms. MW: mind-wandering episodes; 2S and 5S: the last 2 or 5 seconds prior to self-report; FP: the full reading page duration. Asterisks indicate significance levels from permutation tests (n = 200): * p < .05 and ** p < .01. (B) Permutation-based feature importance for the classifier trained on MW episodes (self-report paradigm). Error bars represent 95% confidence intervals computed across LOSO cross-validation folds.

In the simulated thought-probe condition, the classifier trained on the MW-onset window again performed best (AUROC = 0.621), significantly outperforming the 2-second window (AUROC = 0.544, *p* = .010), the 5-second window (AUROC = 0.551, *p* = .015), and the full page window (AUROC = 0.569, *p* = 0.035).

Using logistic regression allowed us to examine feature importance and interpret the contribution of individual features to the final model. As shown in Figure 4B, permutation-based feature importance permutation tests (n = 200): * ***p*** < .05 and ** ***p*** < .01. (B) Permutation-based feature importance for the classifier trained on MW episodes (self-report paradigm). Error bars represent 95% confidence intervals computed across LOSO cross-validation folds.

### Temporal Dynamics of MW Classification Performance

Our task paradigm allowed for precise identification of MW onset, enabling a time-resolved analysis of classifier performance. Using a 2-second sliding window stepped every 0.25 seconds, we extracted eye-tracking features around MW onset and self-report and trained a logistic regression classifier at each time point (Figure 5A). Prior to MW onset, classifier performance hovered around chance (AUROC ≈ 0.5). After MW onset, AUROC increased and peaked at 0.65 around 3 seconds post-onset. When trials were time-locked to the moment of self-report, AUROC stabilized around 0.6 beginning about 5 seconds prior.

**Figure 5.**
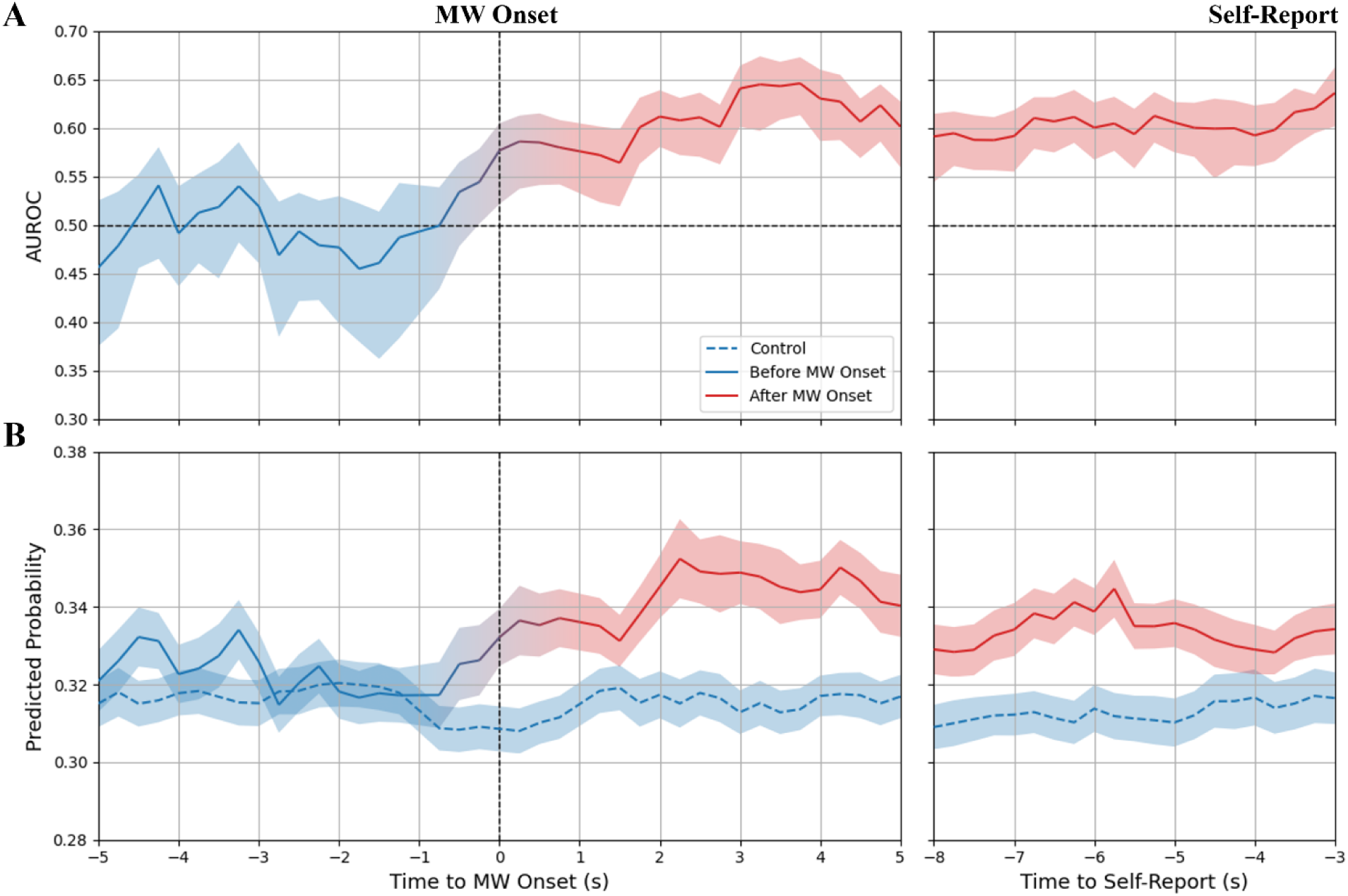
Logistic regression classifier performance at times surrounding MW onset and self-report. *Note.* (A) AUROC score for classifiers trained on data from a 2-second sliding window centered at each time point relative to MW onset (left) and self-report (right). Blue lines are outside of a MW episode, and red lines are during MW. The color gradient from blue to red ranges from –1 s to 1 s relative to MW onset, which corresponds to the 2-second sliding window used for feature extraction. Shaded areas represent a 95% confidence interval based on 200 permutations. The horizontal line at AUROC = 0.5 represents the AUROC expected by random chance. (B) Average probability of MW predicted by a classifier trained on data at MW onset (0 s) and applied to features at the time point on the x axis, with separate lines averaging across MW trials (solid line) and control trials (dashed line). Shaded areas represent the standard error across 44 participants.

To test whether the discriminative power of features remained stable during MW, we applied the classifier weights trained at MW onset (0 s) to features from other time points. The resulting predicted probabilities are shown in Figure 5B. Because the same dataset was used for both training and evaluation, the probability at MW onset (0 s) was interpolated to reduce potential overfitting. No significant difference in prediction strength was observed between MW onset (0 s) and the middle of the episode (5 s) (Wilcoxon signed-rank test: *p* = 0.209), suggesting that MW onset did not have a distinct eye tracking signature that was not also present within the MW episode.

### Divergence of Eye Features Following MW Onset

We also examined the temporal dynamics of individual eye-tracking features around MW onset using the same sliding window analysis (Figure 6). For each participant, fixation rate, fixation dispersion, and pupil size were averaged across all pages and runs. No other eye-tracking features showed consistent and stable divergence around MW onset compared to control periods (see Supplementary Figure 7).

**Figure 6.**
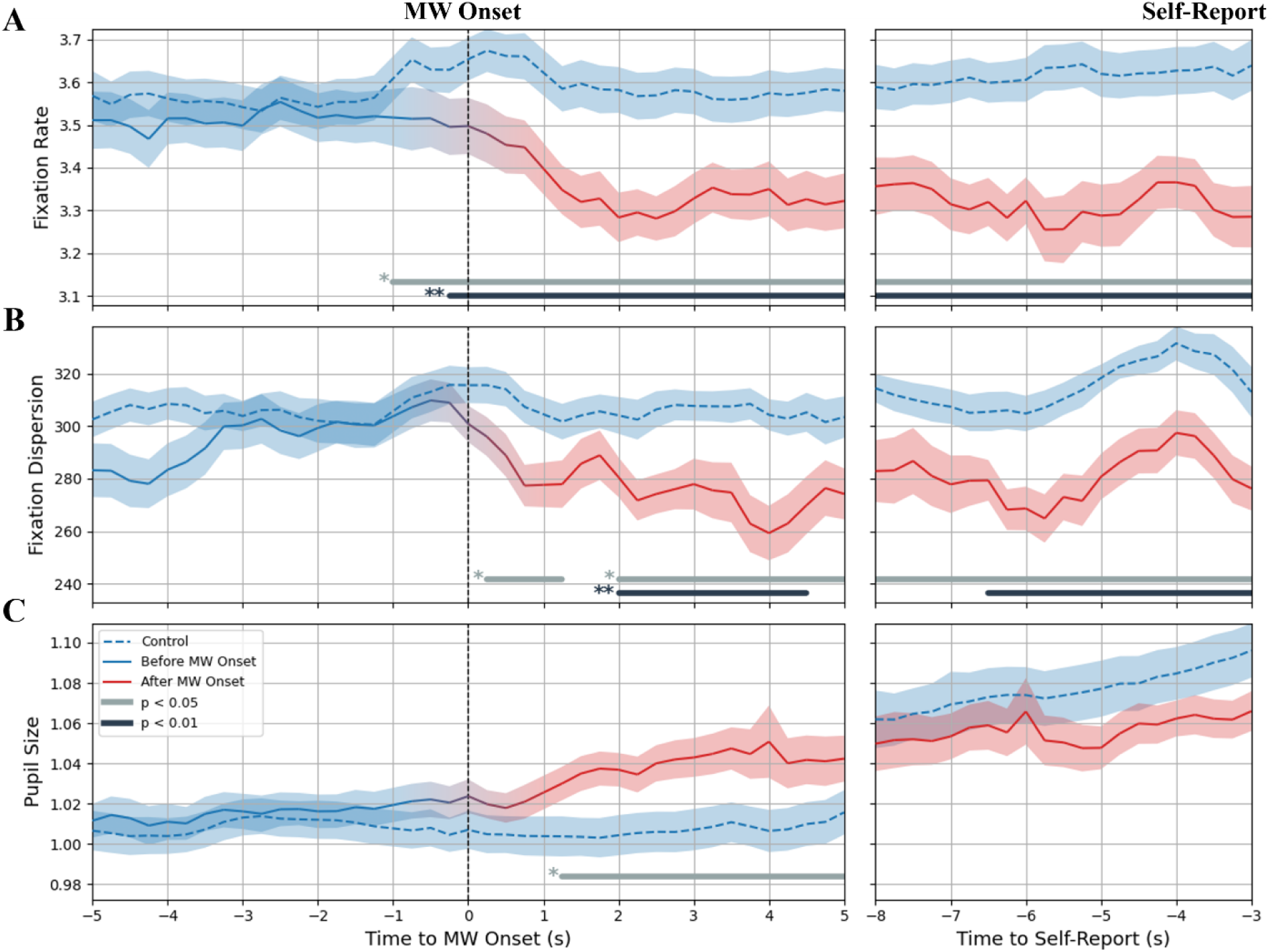
Eye features at times surrounding MW onset and self-report. *Note*. (A) Fixation rate, number of fixations per second. (B). Fixation dispersion, average distance of each fixation from fixation center. (C). Pupil size, normalized by each participant’s baseline (median pupil size across all reading pages). Shaded areas represent the standard error across 44 participants. Asterisks and horizontal bars indicate time intervals with statistically significant differences between control and MW (Wilcoxon signed-rank test, * *p* < .05 and ** *p* < .01, false discovery rate corrected). To reduce false positives from transient fluctuations, only significant intervals lasting longer than 1 s were highlighted and plotted.

In Figure 6A, participants made approximately 3.6 fixations per second during control periods. A small bump from –1 to +1 seconds in the control trace reflects the trial alignment procedure: trials were time-locked to a randomly selected fixation, meaning a fixation is always present at time zero. To improve visualization and capture general trends, fixation rates at exactly -1 and +1 seconds relative to MW and control period onset times were interpolated. Notably, fixation rate began to decline about 0.25 seconds prior to MW onset and continued to drop throughout the MW episode (mean Cohen’s |d| = 0.839, SD = 0.220), eventually reaching about 3.3 fixations per second. This reduction remained statistically significant throughout the entire MW period (*p* < .01, FDR-corrected; mean Cohen’s |d| = 0.973, SD = 0.161).

Similarly, fixation dispersion remained relatively stable during control periods at around 300-pixel units. As shown in Figure 6B, fixation dispersion started to decrease immediately after self-reported MW onset (p < .05, FDR-corrected; mean Cohen’s |d| = 0.515, SD = 0.129). This decrease stabilized about 2 s after MW onset (p < .01, FDR-corrected; mean Cohen’s |d| = 0.592, SD = 0.115), reaching approximately 280-pixel units.

As shown in Figure 6C, pupil size remained stable at baseline and began to rise about 1 second after to MW onset (*p* < .05, FDR-corrected; mean Cohen’s |d| = 0.433, SD = 0.047). Unlike fixation rate and dispersion, pupil size in control trials was not consistent across time. Specifically, pupil size increased from approximately 1.06 to 1.10 in the final 5 seconds before the end of the page. We attribute this rise to increased arousal or anticipatory effects near the end of the reading page (see Supplementary Figure 8).

Because of this shift during control trials, we did not observe a statistically significant difference in pupil size between MW and control conditions when time-locked to the self-report or end of page, despite the increase in pupil size as self-report approached.

To confirm that the pupil dilation following MW onset was not simply due to proximity to the end of the page, we conducted a sensitivity analysis (Supplementary Figure 9). Pupil size consistently increased after MW onset regardless of MW episode duration, supporting the interpretation that the change reflects MW-related cognitive processes rather than page-related arousal.

## Discussion

In this study, we introduced a novel paradigm called ReMind that uses retrospective self-report measures to investigate mind-wandering during reading. Unlike earlier studies that focused on meta-awareness or the end of MW episodes, our approach targets the onset of MW, which is a critical yet largely unresolved aspect of mind-wandering research. A potential concern in interpreting our results is whether participants can accurately report when their MW began. While MW is often characterized as an unconscious or drifting mental state (Schooler, 2002; Shepherd, 2019; Zedelius et al., 2018), several lines of evidence suggest that participants can retrospectively identify MW onset with reasonable accuracy.

First, participants in our task received explicit task instructions, a clear definition of MW, and practice in reflecting on the flow of their attention in normal reading and marking both the onset and offset of MW. Second, our results showed that logistic regression classifier benefited from access to reported MW onset in both self-report and thought-probe paradigms, suggesting that this information better defines the temporal boundaries of MW episodes and allows eye-tracking features to be extracted from more distinct attentional states. Finally, our temporal analyses showed that MW classification performance began rising from chance level around the reported onset and peaked approximately 3 s later. At the feature level, we observed decreases in fixation rate and fixation dispersion, along with increases in pupil size, following the estimated onset across trials and participants. This alignment between subjective reports and objective eye-tracking measures further supports the validity of the reported MW onset. Together, these findings demonstrate the feasibility of identifying MW onset through retrospective self-report and highlight the broader utility of our paradigm for studying the temporal dynamics of attention.

Building on this validation, we examined the frequency and duration of MW episodes. Previous studies have suggested that MW frequency is influenced by both task content (Feng et al., 2013; Kane et al., 2007; Soemer & Schiefele, 2019) and task duration (Krimsky et al., 2017; Zanesco et al., 2025).

Accordingly, we used a generalized linear mixed-effects model to examine these effects in our dataset. To control task content, we selected the reading materials carefully to cover a broad range of topics while maintaining relatively similar levels of engagement across stories. Only one story (Pluto) showed a significant effect on MW frequency, suggesting that the stories were generally well matched overall while still allowing some topic-related variability in MW occurrence. To examine task duration effects, we analyzed both within-story progression and overall experimental progression. Our results showed that the probability of reporting MW increased as participants progressed through an article. However, no significant effects of run order or overall task duration (page order × run order interaction) were observed, suggesting that MW did not systematically increase across the overall experimental session. Therefore, the page effect is more likely related to the unfolding content of each story, because page progression was inherently coupled with story progression. In other words, later sections of the stories may have contained content that was more likely to induce MW. Together, these findings suggest that the present task reasonably controlled for broad effects of task content and duration. At the same time, substantial participant-level variance estimated both by the model and by individual MW frequencies suggests strong individual differences in the tendency to report MW. This variability further highlights both the difficulty and importance of accounting for participant-level differences when studying the temporal dynamics and behavioral signatures of MW.

Across 1,001 reported mind-wandering episodes, the median duration was 6.14 seconds, and the mean was 7.99 seconds. These numbers are comparable to those reported by Pelagatti et al., (2020) who found a median duration of 6.19 seconds and a mean of 8.64 seconds in a cued vigilance task. Like this previous study, we observed a wide range in MW duration: many episodes were very brief, but several lasted up to a minute. Although their paradigm involved cued mind-wandering triggered by specific words, the similarity in duration distributions suggests a potentially shared cognitive mechanism that governs the transition back to task-focused attention. This convergence implies that, regardless of whether mind-wandering is spontaneous or externally cued, the timing of attentional re-engagement may follow a common temporal pattern.

To evaluate how temporal windowing affects MW detection using eye-tracking features, we compared our approach to commonly used fixed-window methods from prior studies. Our results showed that incorporating precise information about MW onset improved classifier performance, particularly relative to full-page analysis methods in both self-report and thought-probe paradigms. The reduced accuracy of full-page classifiers can be attributed to the inclusion of substantial on-task reading time prior to MW onset. In our data, the median interval between page onset and MW report was 18.90 seconds, meaning much of the full-page window contained features associated with focused reading. This likely diluted MW-related signals and impaired classification. In addition, in the simulated thought-probe condition, classifiers trained with known MW onset significantly outperformed those trained on fixed 2-or 5-second window. This likely reflects the limitations of using fixed windows in thought probes, as they rely on participant reflection following an external cue, rather than internally triggered meta-awareness (Zedelius et al., 2015). In addition, thought probes often interrupt MW episodes prematurely, leading to shorter durations and less data from which to compute features. Thus, many 2S or 5S examples in the MW class might include features derived in part from the focused reading periods that precede MW onset. The random timing of thought probes introduces greater uncertainty at MW offset, reducing the reliability of features aligned to the report moment. In contrast, using the self-report moment at the end point, classifiers trained on features extracted from self-reported MW windows did not significantly outperform those using the last 2- or 5-second windows. This may be due to the relatively short MW durations in our dataset, which were often brief enough that fixed windows captured most of the relevant signal. Since the self-reported windows may have overlapped substantially with the fixed windows, eye-tracking features extracted from both types of windows are likely to reflect similar cognitive states, leading to the comparable classifier performance.

Overall, these machine learning results quantify the importance of precise temporal alignment in studying cognitive and physiological markers of MW. While a self-reported MW window in principle has the advantage of including more relevant data in long MW episodes and excluding irrelevant data from short episodes, in practice, a fixed 2- or 5-second window prior to a self-report is sufficient to reveal the combination of gaze features that are most predictive of MW. However, in a thought probe paradigm, this does not appear to be the case, and knowledge of MW onset times provides added predictive value.

Given our paradigm’s ability to uniquely localize MW onset, we aimed to examine how MW unfolds over time. We applied a sliding-window approach to extract features at fine-grained temporal intervals, aligning trials to either MW onset or self-report. To assess the dynamic separability of MW and on-task states, we trained classifiers at each time point. Our results revealed a gradual rise in classifier performance starting before MW onset and peaking during the stable MW phase. To our knowledge, this is the first study to evaluate MW classifier performance near onset with such temporal resolution. In contrast to earlier work that relied on broad or ambiguous windows (D’Mello et al., 2020; Faber et al., 2018; Mills et al., 2021; Reichle et al., 2010), our findings have implications for real-time MW detection systems, particularly for enabling timely interventions. We also find that a classifier trained to predict MW at one point (e.g., MW onset) works similarly well at other points during MW, an encouraging sign for sliding-window classifiers and real-time interventions.

To gain insight into the specific features driving classification, we examined feature importance in classifiers trained on self-reported MW episodes and plotted their temporal dynamics. Among all features, fixation rate emerged as the most informative. The time series revealed a marked decline in fixation rate beginning near MW onset and stabilizing thereafter. This aligns with prior findings that mindless reading is characterized by reduced fixation rates and longer fixation durations (Foulsham et al., 2013; Reichle et al., 2010). Similarly, fixation dispersion decreased following MW onset. Because fixation dispersion was calculated as the root mean square distance between individual fixations and their average spatial position, lower values indicate that fixations became more spatially concentrated during MW episodes. One possible interpretation is that readers continued making eye movements across the text, but with reduced spatial exploration once attention disengaged from comprehension. Interestingly, this pattern differs from one previous study reporting increased fixation dispersion during MW (Faber et al., 2020), possibly due to differences between sentence-level reading tasks and our more naturalistic page-reading paradigm. Whereas the previous study presented only a few sentences at a time, our task required continuous scanning across 16 full lines of text, such that MW may lead to a narrowing rather than broadening of gaze distribution. Because reading requires coordinated fixations and saccades to support information processing (Dambacher et al., 2013; Rayner et al., 2012), fixation rate and fixation dispersion may serve as behavioral proxies for attentional allocation, reflecting how rapidly and broadly readers sample information from the text (Ban et al., 2024). Our results provide further support for the perceptual decoupling hypothesis (Smallwood & Schooler, 2006), which posits that internal thoughts during MW divert resources away from external stimuli. In addition, they suggest that this observable change is immediate and stable rather than increasing with MW duration.

The consistent alignment between subjective reports and objective eye-tracking changes suggests that the reported onset corresponds to a reliable transition in attentional state. However, one possible interpretation of our findings is that this transition reflects the point at which MW becomes behaviorally detectable or begins to disrupt comprehension or memory for the text, rather than the earliest moment at which MW emerges. This interpretation is consistent with our temporal analyses showing that classifier performance continued to increase and certain features started to diverge after the reported onset.

However, if the earliest stages of MW occur unconsciously and do not yet influence reading behavior, comprehension, or awareness, then such states may have limited behavioral relevance and may not be accessible through retrospective reports or any existing method relying on behavioral or subjective ground-truth labels. From this perspective, the transitional state captured by ReMind corresponds to the point at which eye-tracking features begin to show noticeable differences and MW starts to meaningfully interfere with ongoing task-related thought, which we would argue is the most behaviorally relevant aspect of MW onset during reading.

We also examined features related to reading content, such as regressions and lexical variables (e.g., word frequency), which have been shown to influence MW classification in previous studies (Foulsham et al., 2013; Reichle et al., 2010; Schad et al., 2012). However, in our sliding-window analysis, these features generally did not exhibit consistent divergence from control periods following MW onset, with the exception of a brief increase in in-word regression values (see Supplementary Figure 7). This may be due to their sensitivity to window duration. In our time series, features were calculated using 2-second windows that may contain too few data points to compute stable estimates, especially for metrics involving correlation (e.g., frequency-fixation relationships). Consequently, these features were noisier and contributed less to classification in this analysis, even though they may be more reliable when computed over longer durations.

Pupil size has been widely studied in relation with cognitive state and arousal (Castellotti et al., 2025; Joshi & Gold, 2020; Pelagatti et al., 2025; Wang et al., 2018), However, the literature on pupil dynamics during MW is mixed, with studies reporting increases (Franklin et al., 2013; Groot et al., 2021; Oyarzo et al., 2022; Pelagatti et al., 2020; Smallwood et al., 2011), decreases (Gouraud et al., 2018; Grandchamp et al., 2014; Mittner et al., 2014; Ozawa et al., 2022), or no significant differences (Groot et al., 2022; Hood et al., 2022; Uzzaman & Joordens, 2011) in pupil size depending on the specific task context. In our time-series plots, we observed a clear increase in pupil size following MW onset. At the same time, control trials time-locked to page endings also showed elevated pupil size in both MW and non-MW conditions. This suggests a page-level effect, possibly linked to motor preparation (e.g., finger movement to advance pages) or implicit tracking of task progress (Oyarzo et al., 2022). While we haven’t identified the exact cause, this effect likely explains why pupil size ranked low in feature importance when training classifiers on all MW episode data. Importantly, we confirmed that the pupil size increase near MW onset is not an artifact of trials being closer to page end by stratifying trials by MW duration. The effect persisted across duration bins, supporting its link to MW rather than task timing.

Together, these dynamic feature trajectories offer novel insights into theoretical models of MW. The fixation rate and dispersion decline supports the perceptual decoupling hypothesis by marking reduced perceptual engagement starting at MW onset. We also evaluated predictions from the “Neural Model of Mind-Wandering” proposed by Mittner et al., (2016), which suggests that elevated tonic norepinephrine activity induces an exploratory state, marked by pupil dilation. We did not observe the peak in pupil size around MW onset that such a model might suggest (although this off-focus state can also occur within MW or on-task periods). Moreover, the feature set that was predictive at MW onset did not become less predictive when applied to windows during the middle of MW episodes, providing little support for a unique transitional “off-focus” state.

Finally, our findings have implications for future MW detection systems. Many prior classifiers are limited by paradigms that lack precise temporal information about MW onset, making it difficult to anchor feature extraction to meaningful cognitive events. Our time-resolved feature trajectories offer a data-driven benchmark for identifying the onset of MW. Classifiers trained on other datasets could potentially incorporate these feature trajectories as a template or ground truth, enabling them to infer MW onset without requiring new data collection. This approach could guide the development of more temporally precise and cognitively informed models for detecting the onset and dynamics of mind-wandering.

Our approach has limitations. Because recruitment and data collection were conducted at a university, our sample was dominated by younger adults (median = 20, mean = 22.6), which may limit the generalizability of our findings to broader populations. Reading behavior (Rayner et al., 2006), mind-wandering frequency and duration (Giambra, 1989b), and willingness to self-report MW (Jackson & Balota, 2012; Seli et al., 2021) may vary with age. As a result, the temporal dynamics and eye-tracking patterns associated with MW observed in the present study may differ in older adults or more diverse populations. Future work should therefore examine the robustness of ReMind across a wider range of demographic and cognitive backgrounds.

We selected a self-report rather than thought-probe approach because our goal was to combine retrospective reports with continuous reading behavior to estimate MW onset while preserving the natural flow of reading and avoiding interruptions to ongoing cognitive processing. However, self-caught approaches rely heavily on meta-awareness and are likely to miss MW episodes that are brief, weak, or never become consciously noticeable (Sayette et al., 2009). In contrast, thought-probe paradigms may sample a broader range of MW states, including episodes that participants would not spontaneously report, although this comes at the cost of poorer temporal precision and disruption of ongoing reading behavior (Kane et al., 2021). As a result, the present paradigm likely captures a subset of MW episodes that become sufficiently salient to trigger awareness and self-report. This may influence both the frequency of detected MW episodes and the eye-tracking patterns learned by the classifier. Nevertheless, for the present study, which aimed to estimate the temporal dynamics surrounding MW onset during naturalistic reading, we believe the self-report approach was the most appropriate design choice. Future work may benefit from combining this retrospective idea with thought-probe paradigms to better capture a broader spectrum of MW experiences depending on the goals of the study.

MW episode durations varied substantially across trials, ranging from very brief episodes to a smaller number of relatively long ones. Although we interpret these episodes within the framework of mind-wandering, some longer episodes may also involve more sustained or constructive internally directed thought, such as autobiographical planning or memory-related processes. In the present study, our temporal analyses aligned trials to two relatively stable reference points: the retrospectively reported MW onset and the self-report moment. This approach was designed to capture the general transition into and continuation of MW across episodes. However, it is possible that MW episodes with different durations reflect different underlying temporal structures, such that the emergence, maintenance, and disengagement of MW vary across trials. Under these conditions, aligning all episodes solely by onset and self-report may obscure meaningful heterogeneity in MW dynamics. In addition, for simplicity and consistency with prior work, we aligned MW trials to end at the self-report moment rather than participants’ reported MW offset time. This choice helped control for potential confounds related to page-ending effects and ensured fairer comparisons between MW and control trials. However, it may overlook cognitive processes occurring between MW offset and self-report. Future work should therefore examine whether different classes or durations of MW episodes exhibit distinct temporal profiles and should further investigate the transition period between MW offset and self-report.

Although eye-tracking provides a non-invasive and scalable tool for assessing attentional state (Brishtel et al., 2020b; Mills et al., 2021), it remains a relatively coarse proxy for underlying brain processes (Hessels & Hooge, 2019). Commonly used eye-tracking features, such as fixations and saccades, require a time window for computation and cannot provide true moment-to-moment resolution. Even with our sliding-window approach, eye features are temporally smoothed, limiting their sensitivity to rapid changes. Pupil size, while temporally continuous and strongly linked to attention and arousal, is sensitive to physical task conditions, such as luminance and screen layout (Pan et al., 2024). In our reading task, page-start and page-end effects further confounded pupil dynamics. These findings underscore the importance of task design in interpreting physiological signals and suggest that future experiments should aim to minimize such confounds (Sperandio et al., 2018).

To assess generalizability, we used leave-one-subject-out cross-validation, which accounts for individual variability and simulates real-world classifier deployment. Classifier performance was computed by aggregating predictions across all test folds rather than averaging participant-level AUROC scores. In practice, this aggregated approach implicitly weights the overall performance by participant sample size, such that participants contributing more samples have a larger influence on the final AUROC estimate. However, some participants contributed relatively few MW trials, which may still introduce variability and reduce overall performance metrics. We chose not to upsample these sparse cases to avoid introducing bias. A more robust strategy would involve recruiting more participants and applying stricter inclusion criteria to ensure a balanced distribution of MW and non-MW trials per participant. We could also build personalized classifiers tailored to individual differences, but this would require much longer task durations and more extensive data collection. That said, task duration must be carefully managed.

Prior research shows that longer tasks can lead to mood deterioration (Jangraw et al., 2023), which may affect mind-wandering frequency (Smallwood et al., 2009; Welhaf et al., 2024) and reduce ecological validity. Thus, future experimental designs must balance the need for large-scale data with participant comfort and attentional engagement to better reflect real-world conditions.

In future work, we plan to expand our investigation into how linguistic content influences gaze behavior. In the current study, lexical features such as word length and frequency did not significantly contribute to classification when calculated using short sliding windows. One likely limitation is our use of static frequency scores from English corpora, which do not account for semantics and contexts (Piantadosi, 2014). Because words can carry different meanings depending on the surrounding text, their gaze-related effects may be modulated by context. To address this, we intend to incorporate natural language processing techniques, such as contextual embeddings derived from large language models (Kun et al., 2023; Salicchi et al., 2021, 2023), to capture richer semantic representations from the full story context. By correlating these semantic vectors with fixation metrics, we hope to uncover more nuanced relationships between task content and human attention, ultimately improving feature extraction and classification models.

In conclusion, ReMind provides a retrospective self-report framework for estimating the perceived onset of MW during naturalistic reading. Incorporating participants’ estimated onset timing significantly improved classifier performance, highlighting the importance of onset-level temporal resolution for understanding and detecting MW. Among the most informative features was fixation rate, a key indicator of attentional resource allocation during reading. These results provide direct evidence of physiological changes around MW onset and deepen our understanding of how MW begins and unfolds. By advancing both methodological and theoretical frameworks, this work lays the groundwork for future research on the temporal dynamics of internal attention.

## Acknowledgements

We sincerely thank all the participants for their time and involvement in this study. We also appreciate the undergraduate research assistants from the Glass Brain Lab for their help with data collection.

## Declarations

## Ethics approval

The study protocol was approved by the University of Vermont’s Institutional Review Board

## Consent to participate

All participants provided informed consent prior to participating in the study.

## Consent for publication

All authors have consented to publication of this manuscript.

## Availability of data and materials

All data used in this manuscript is available at DOI 10.17605/OSF.IO/AJME7. Task materials are available at https://github.com/GlassBrainLab/ReMind.git.

## Code availability

The code used to create the task and perform all analyses is available at https://github.com/GlassBrainLab/ReMind.git.

---

**Supplementary Figure 1.**
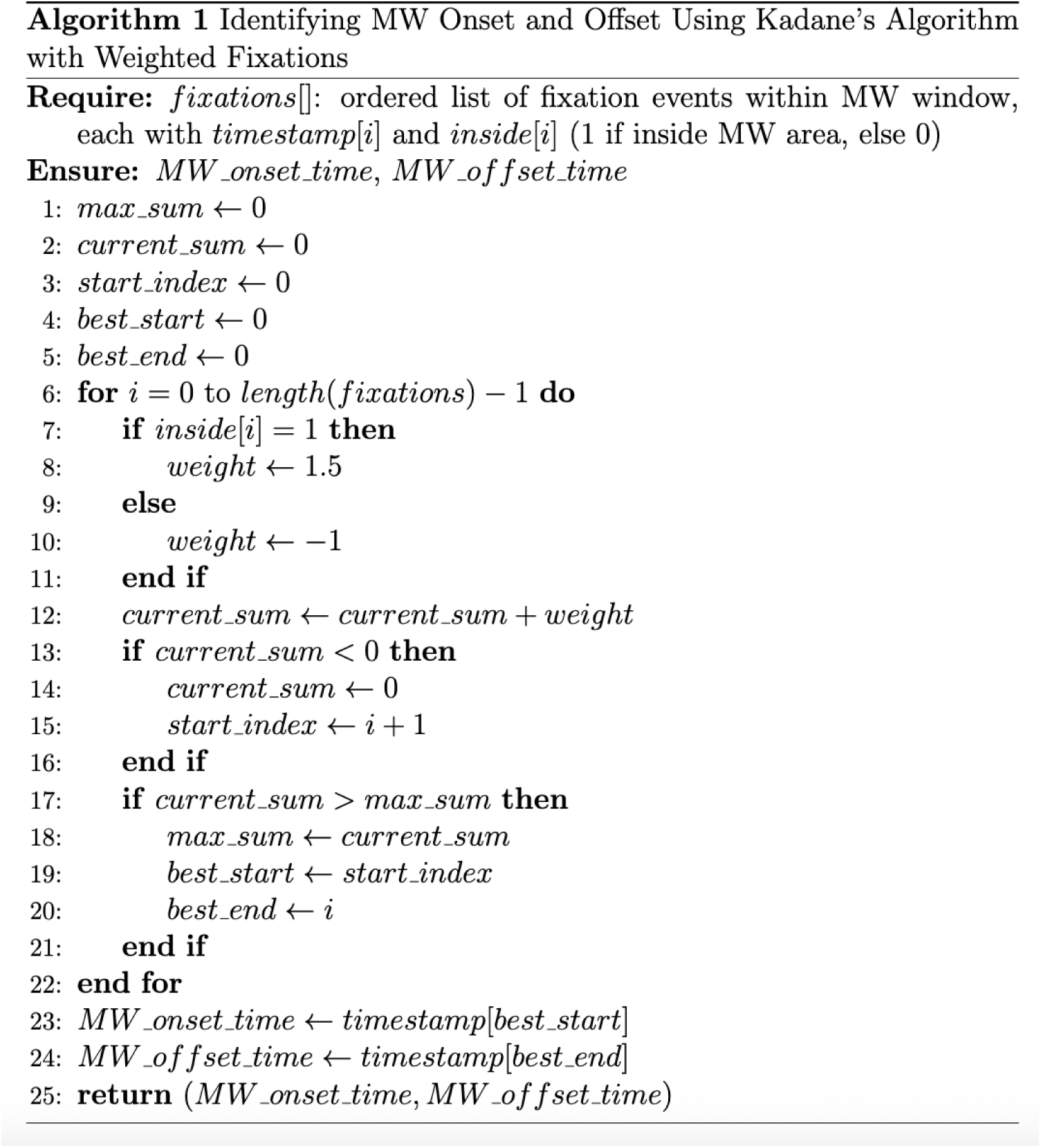
Pseudocode of Kadane’s algorithm for identifying MW onset and offset. *Note*. The algorithm identifies the longest subsequence of fixations within the self-reported words indicating MW onset and offset. It assigns a weight of +1.5 for each fixation inside the MW area and -1 for each fixation outside. This weighted scoring scheme allows the algorithm to tolerate minor deviations outside the reported region while maximizing the density of inside fixations. The start time of first and last fixations from the subsequence with the maximum cumulative weight are selected as the MW onset and offset times, respectively.

**Supplementary Figure 2.**
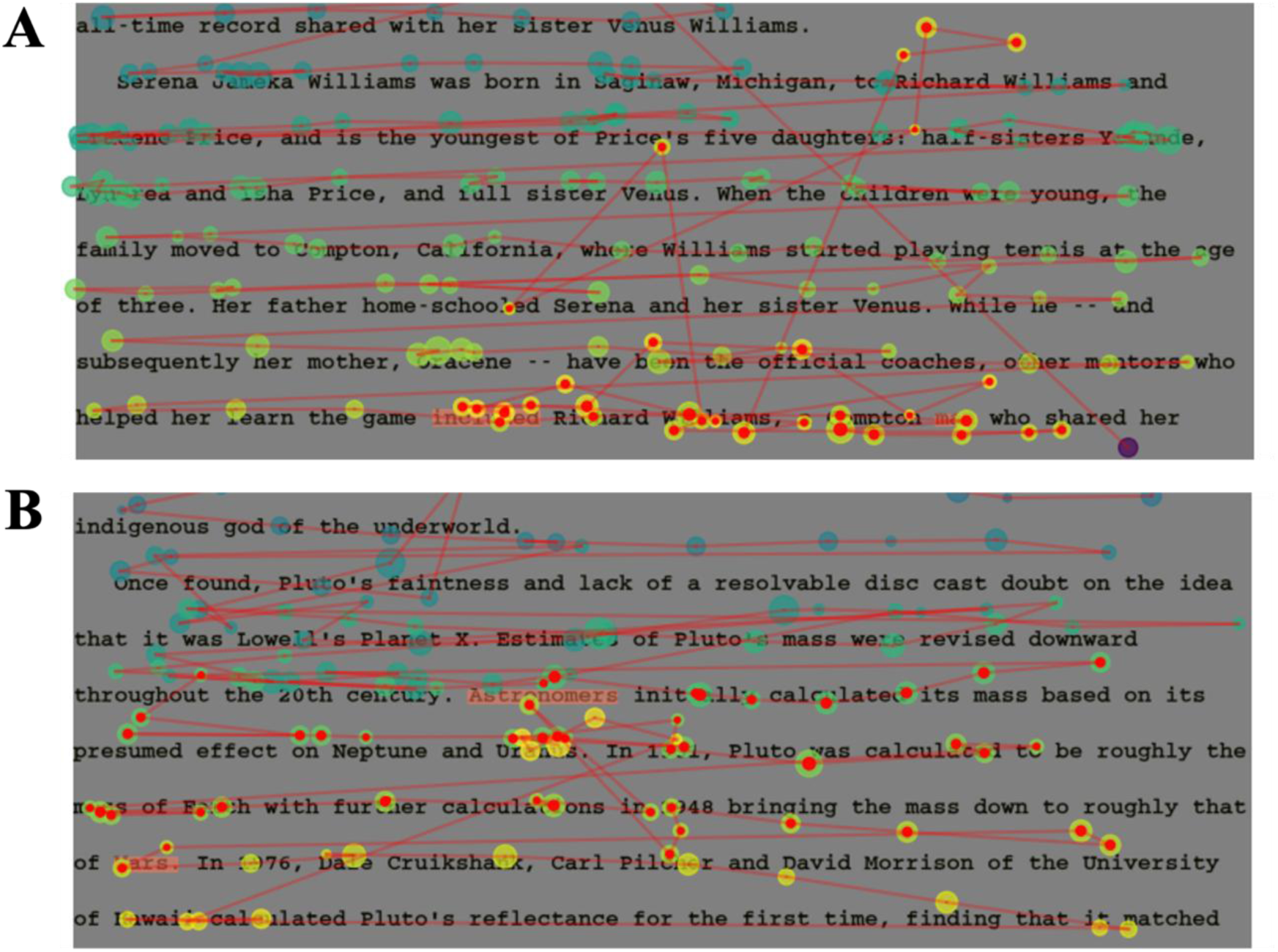
Examples illustrating how Kadane’s algorithm handles continuous fixations that move into and out of the area between two clicked words. *Note*. (A) The participant made fixations outside the designated area. Because the algorithm identifies the longest consecutive sequence of fixations within the area and uses the first and last fixations of that sequence as the MW onset and offset, any fixations outside the area but within the sequence are still labeled as MW fixations. This approach is reasonable because during MW, participants are likely engaged in internal thoughts and less aware of their gaze position or the text being read. (B) An example where fixations after the reported MW offset returned to the area. The algorithm still correctly identifies only the longest consecutive sequence as MW fixations. Even though several fixations after the clicked MW offset fall back into the area, these are not included as MW fixations, preventing an overestimation of MW duration.

**Supplementary Figure 3.**
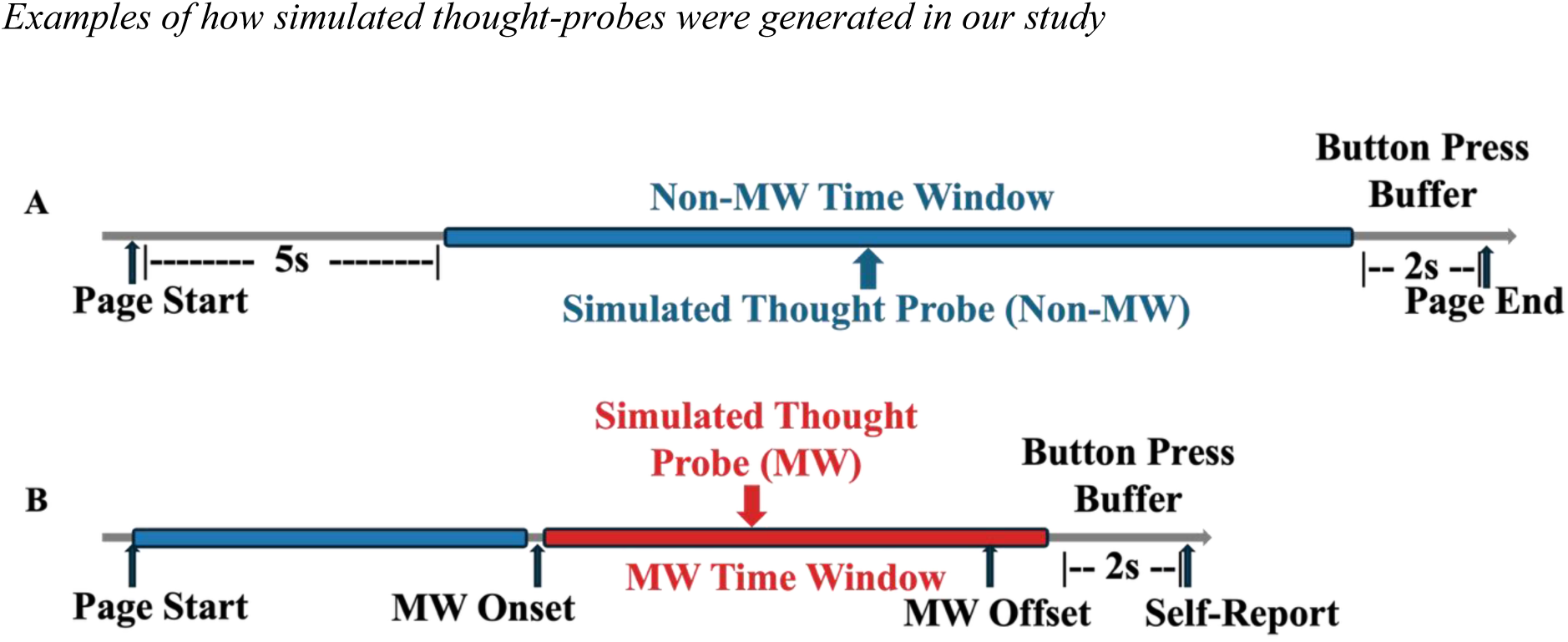
Examples of how simulated thought-probes were generated in our study. *Note.* To avoid confounds, the last 2 seconds before the page ended were excluded. Each simulated thought-probe marked the end of a classifier time window. **(A)** For pages where participants reported no MW, random time points were chosen starting 5 seconds after the page began, since thought-probes typically occur some time after trial onset (here, each page is treated as a trial). These probes were labeled as non-MW. **(B)** For pages reported as MW, random time points were selected between the MW onset and 2 seconds before the self-report button press. These probes were labeled as MW.

**Supplementary Figure 4.**
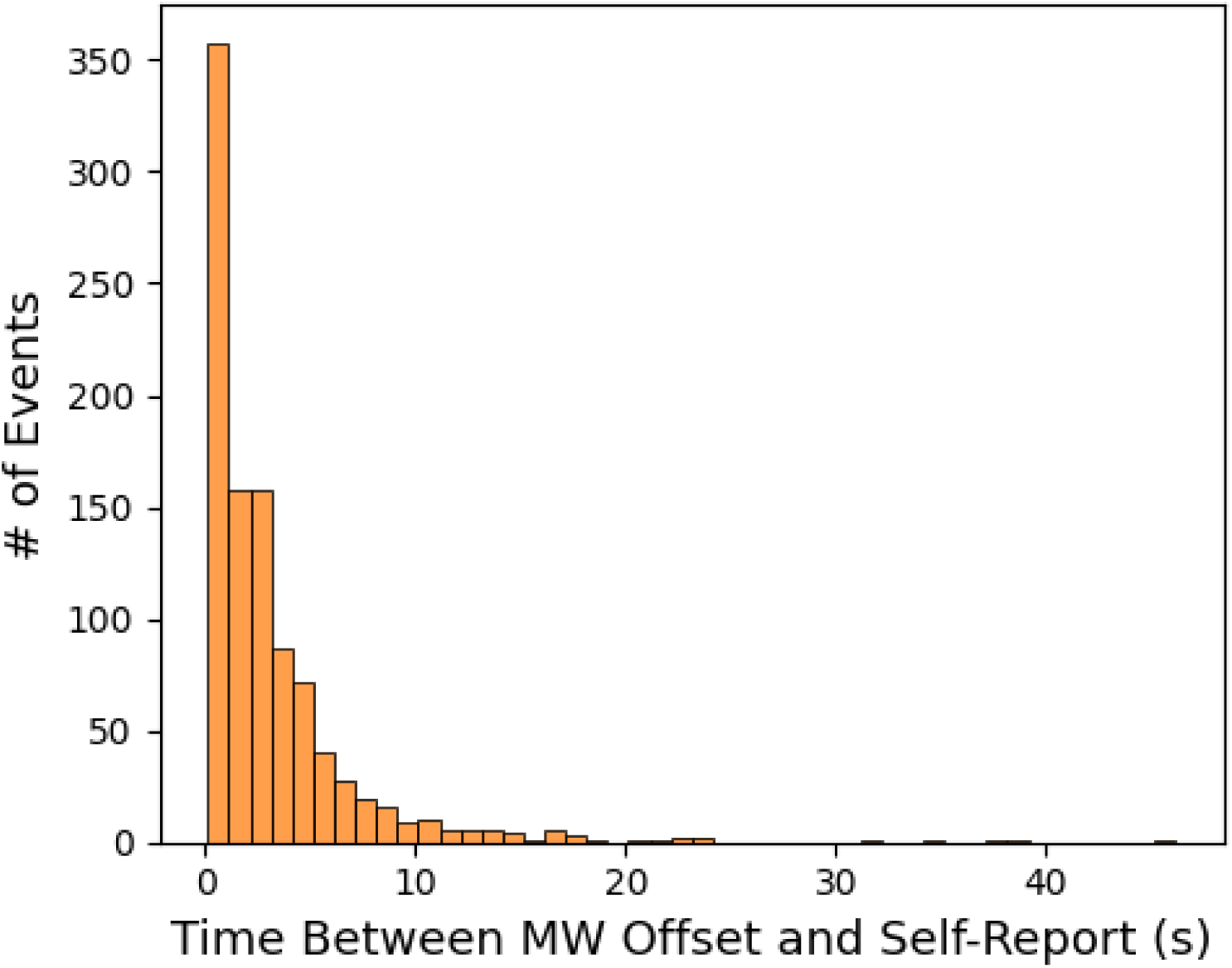
Histogram showing the distribution of time intervals between the offset of MW episodes and the moment of self-report (when they pressed the button) *Note*. On average, MW were reported 3.21 seconds after offset (median = 2.06 seconds; SD = 4.23 seconds). Many MW episodes were reported within a few seconds of their offset, indicating that participants responded promptly after regaining awareness.

**Supplementary Figure 5.**
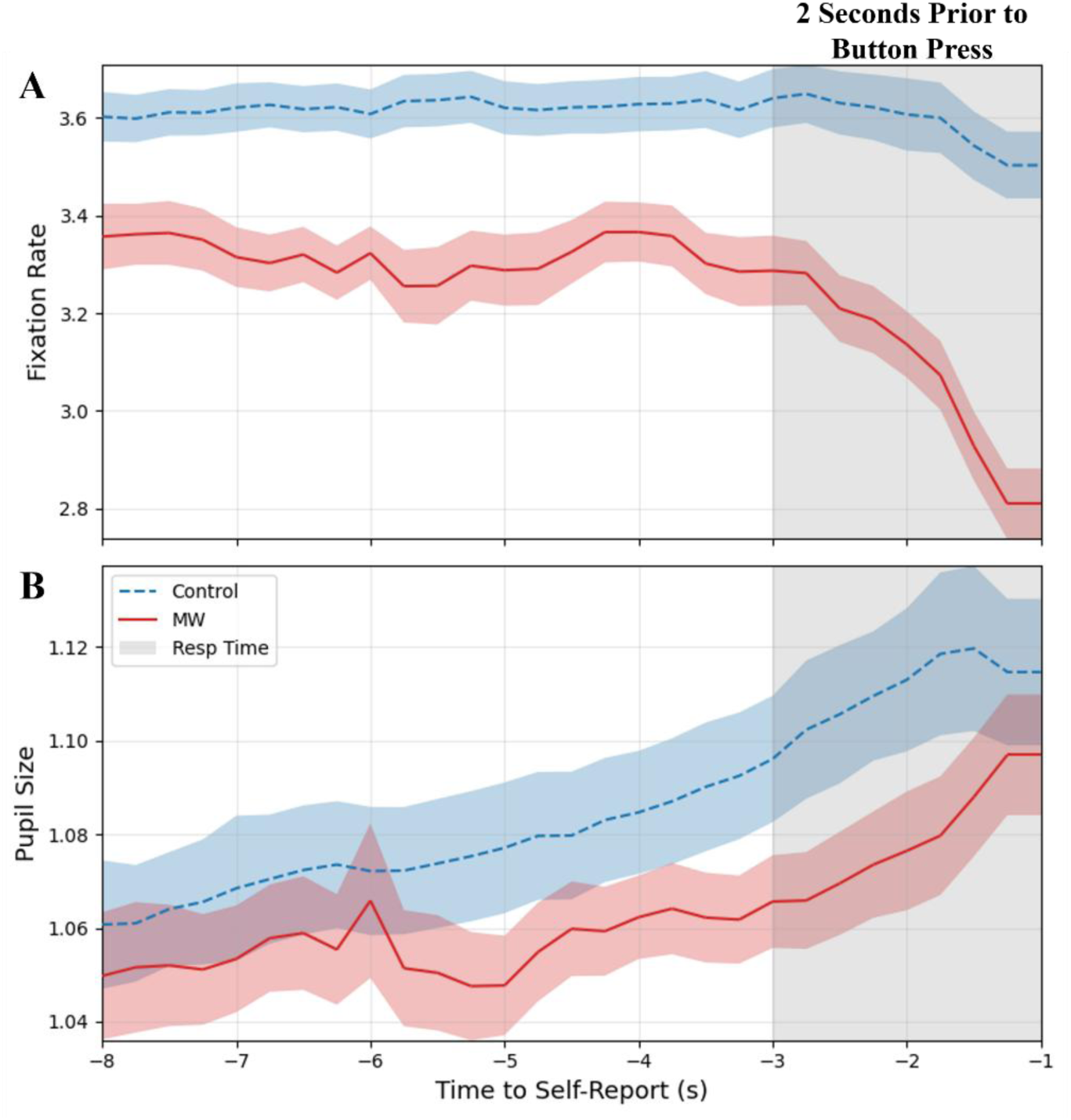
Eye-tracking features extended to the moment of button press. *Note*. Temporal dynamics of (A) fixation rate and (B) pupil size near the moment of button press averaged across 44 participants. Solid red lines represent MW trials and dashed blue lines represent control trials. Shaded regions indicate standard error across participants. Trials were aligned to the moment of button press: participants pressed the “F” key to self-report MW and the “SPACE” key to continue to the next page during control trials. The final 2 seconds before the button press that (based on this figure) we decided might be most strongly affected by motor preparation or arousal are shaded in gray.

**Supplementary Figure 6.**
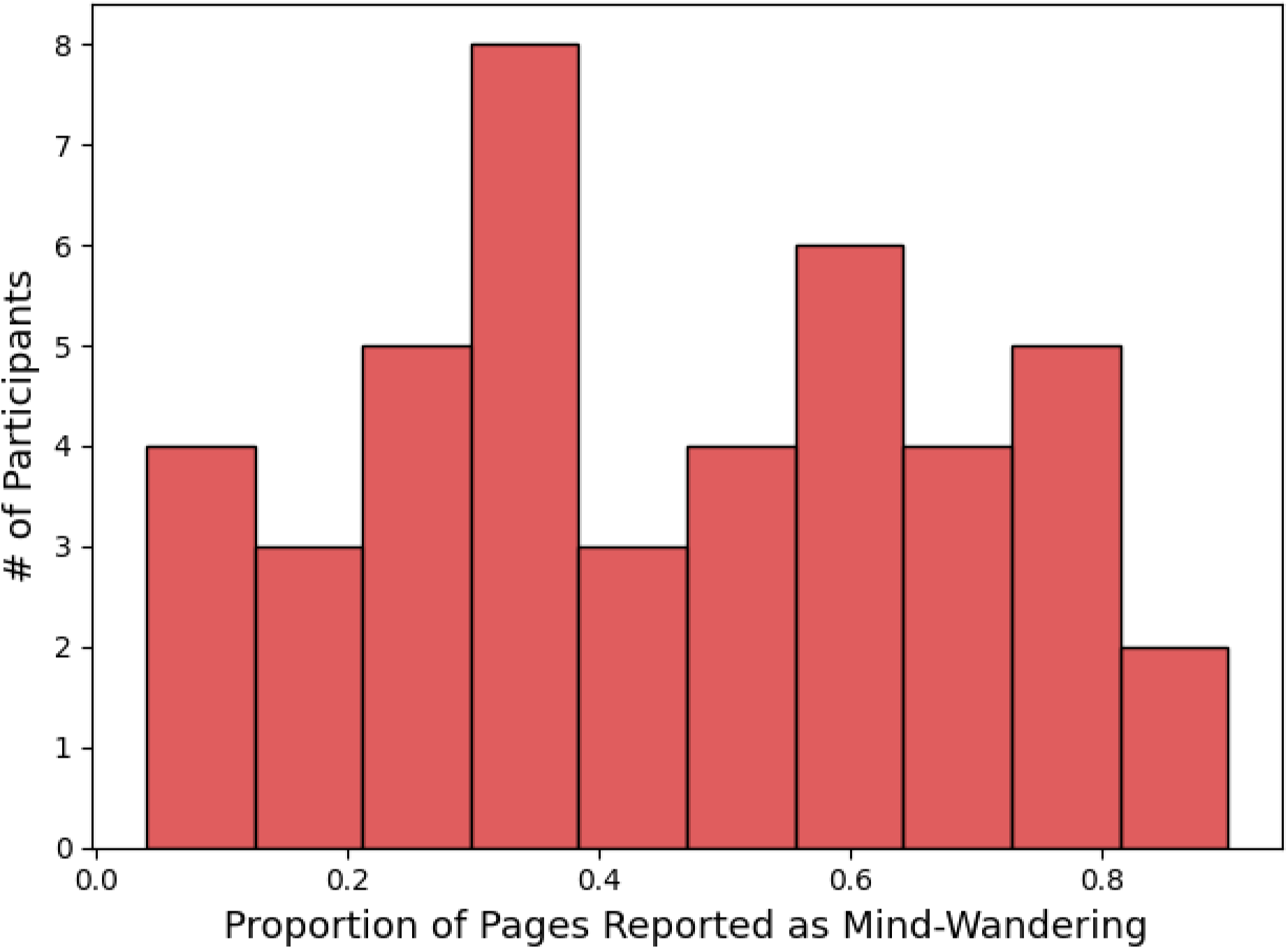
Histogram showing the distribution of individual mind-wandering (MW) proportions across participants. *Note.* Individual MW ratios ranged from 0.04 to 0.90, corresponding to 2 to 45 MW reports out of 50 reading pages. The mean proportion was 0.456 (median = 0.44, SD = 0.236).

**Supplementary Figure 7.**
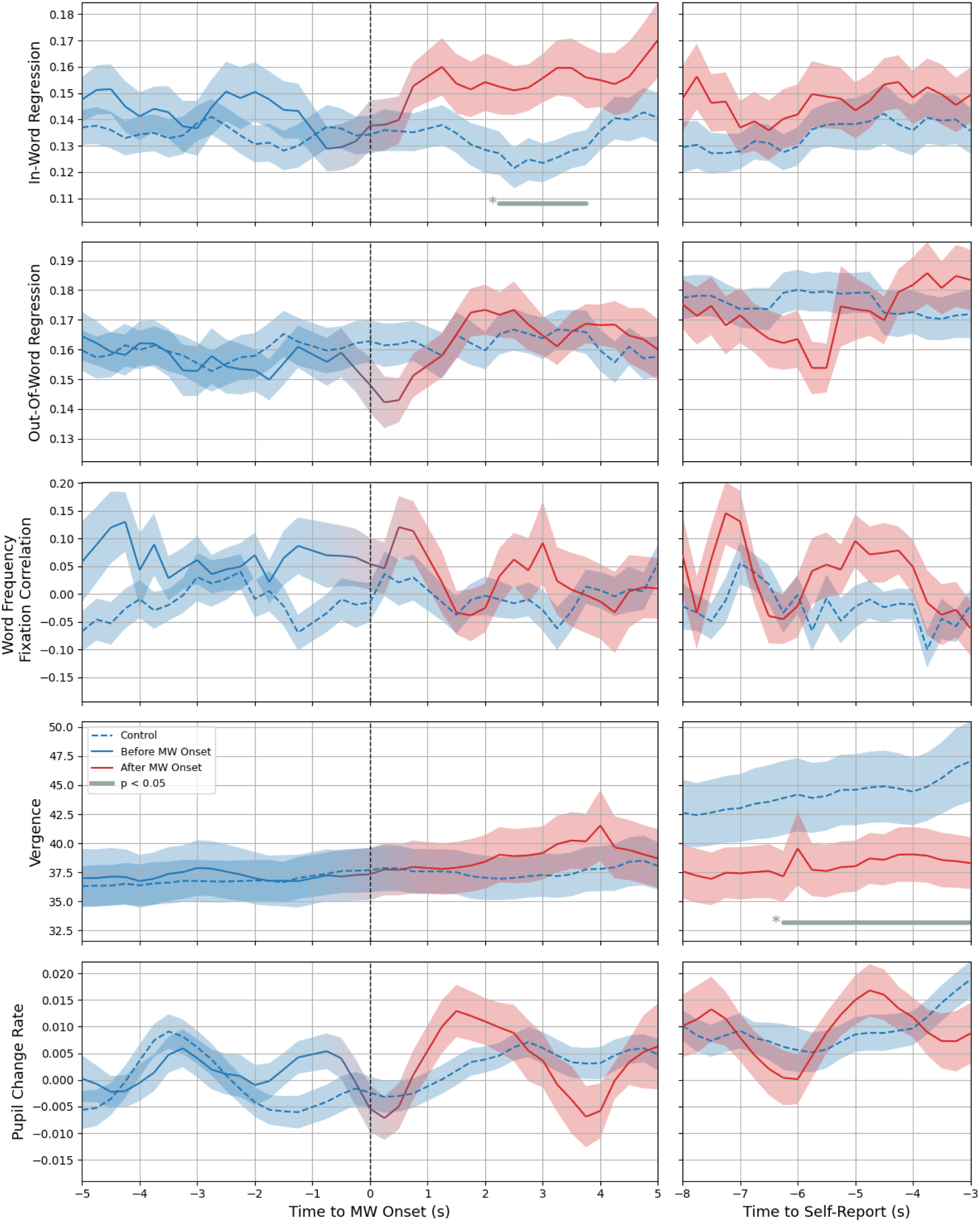
The remaining five eye-tracking features at time points surrounding MW onset and self-report. *Note*. Eye-tracking features were calculated using the sliding-window approach, and the mean of subject means was plotted. Shaded areas represent the standard error across 44 participants. Asterisks and horizontal bars indicate time intervals with statistically significant differences between control and MW (Wilcoxon signed-rank test, * p < .05, false discovery rate corrected). To reduce false positives from transient fluctuations, only significant intervals lasting longer than 1 s were highlighted and plotted. Among the remaining features, in-word regression showed a brief significant increase during MW approximately 2 s after MW onset. This pattern is consistent with prior findings suggesting that readers may temporarily remain fixated on the same word as attention disengages from the text (Mézière et al., 2025). However, the effect lasted for only about 1.5 s and should therefore be interpreted cautiously. Vergence also showed significant differences near the end of the analysis window, which may reflect differences in fixation location on the page. During control trials, participants were typically reading near the bottom of the page, whereas MW reports could occur at any location during reading. No other eye-tracking features showed reliable differences between MW and control conditions.

**Supplementary Figure 8.**
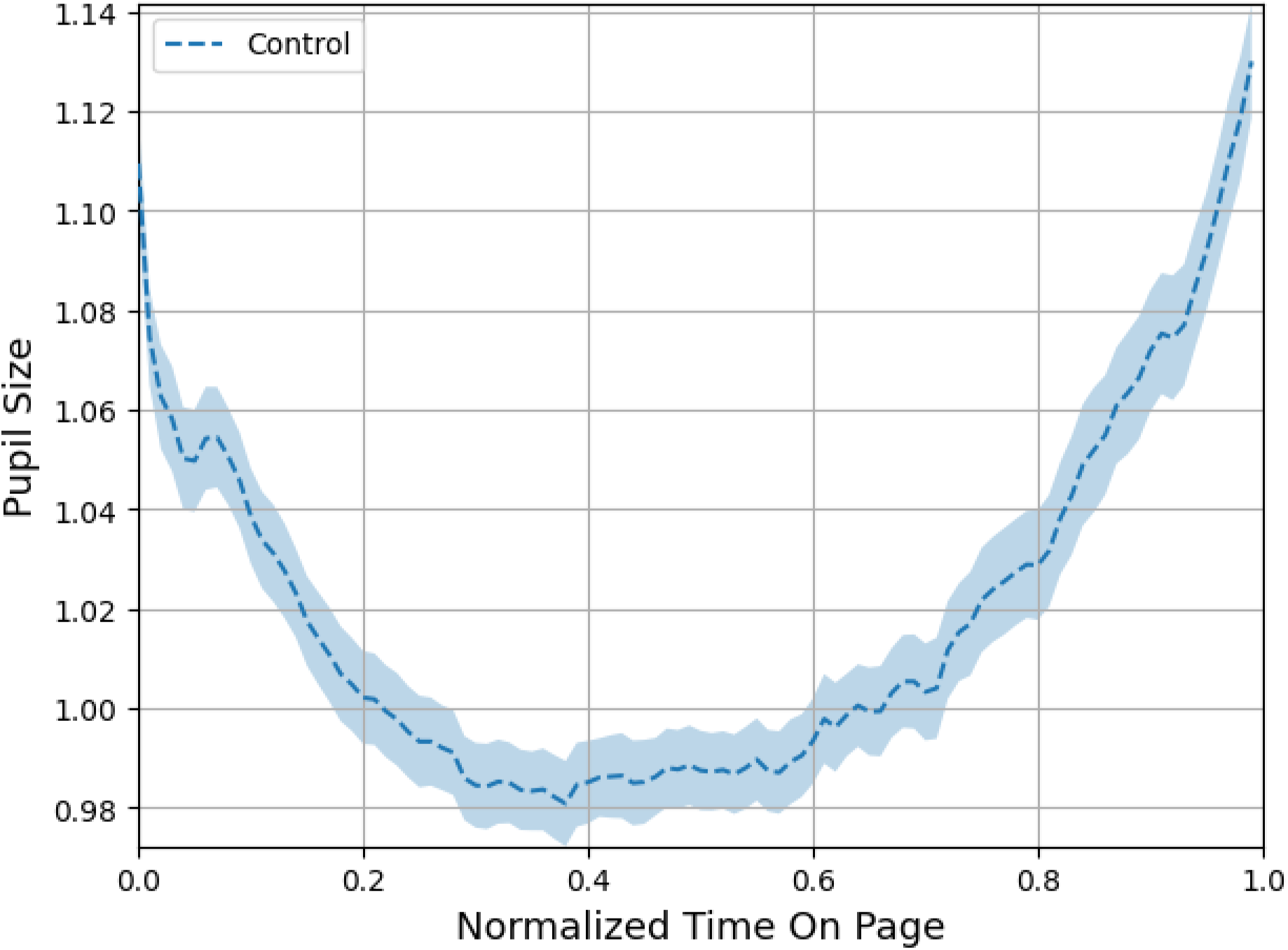
Pupil size changes across normal (non-MW) reading pages. *Note*. Time on page was normalized from 0 (page onset) to 1 (page offset), and only pages without mind-wandering reports were included. The curve reveals a U-shaped pattern, with larger pupil sizes at the beginning and end of the page and a dip in the middle. This effect likely reflects increased arousal or attentional engagement upon entering a new page and anticipatory processing or motor preparation near the end of the page.

**Supplementary Figure 9.**
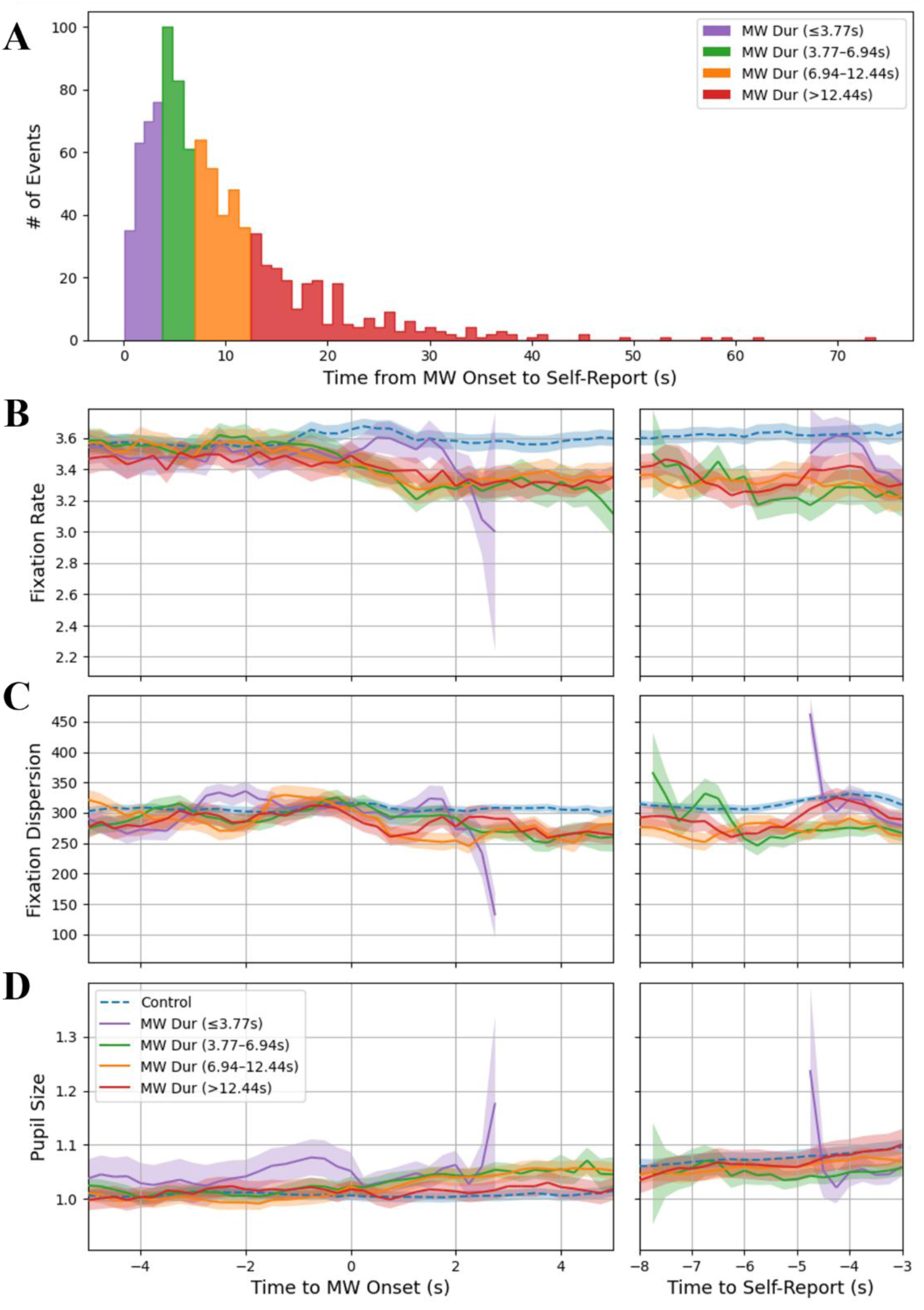
Sensitivity analysis of MW episodes grouped by duration. *Note*. (A) Histogram of MW episodes divided into quartiles based on episode duration, defined as the time from MW onset to 2 seconds before button press. The four duration quartiles are color-coded: purple (≤3.77 s), green (3.77–6.94 s), orange (6.94–12.44 s), and red (>12.44 s). (B) Fixation rate, (C) fixation dispersion, and (D) pupil size time courses surrounding MW onset (left) and self-report (right), plotted separately for each duration group. Control trials (dashed blue) are shown for reference but are not split by duration. Shaded regions represent standard error across participants. Note that the number of trials and participants varies by quartile, as some participants experienced only short or long MW episodes. The purple quartile (shortest duration) shows greater variability near the edges of the analysis window, likely due to limited trial counts, resulting in wider error bands and noisier averages. This analysis demonstrates that the observed changes in fixation rate and pupil size are consistent across MW episodes of varying duration. Specifically, the increase in pupil size observed during MW is unlikely to be driven solely by short episodes occurring near the end of a reading page, where arousal-related effects might occur. Instead, all duration quartiles exhibited comparable trends, suggesting that the feature changes are robust and not driven by any single MW duration subgroup.

**Supplementary Table 1.**
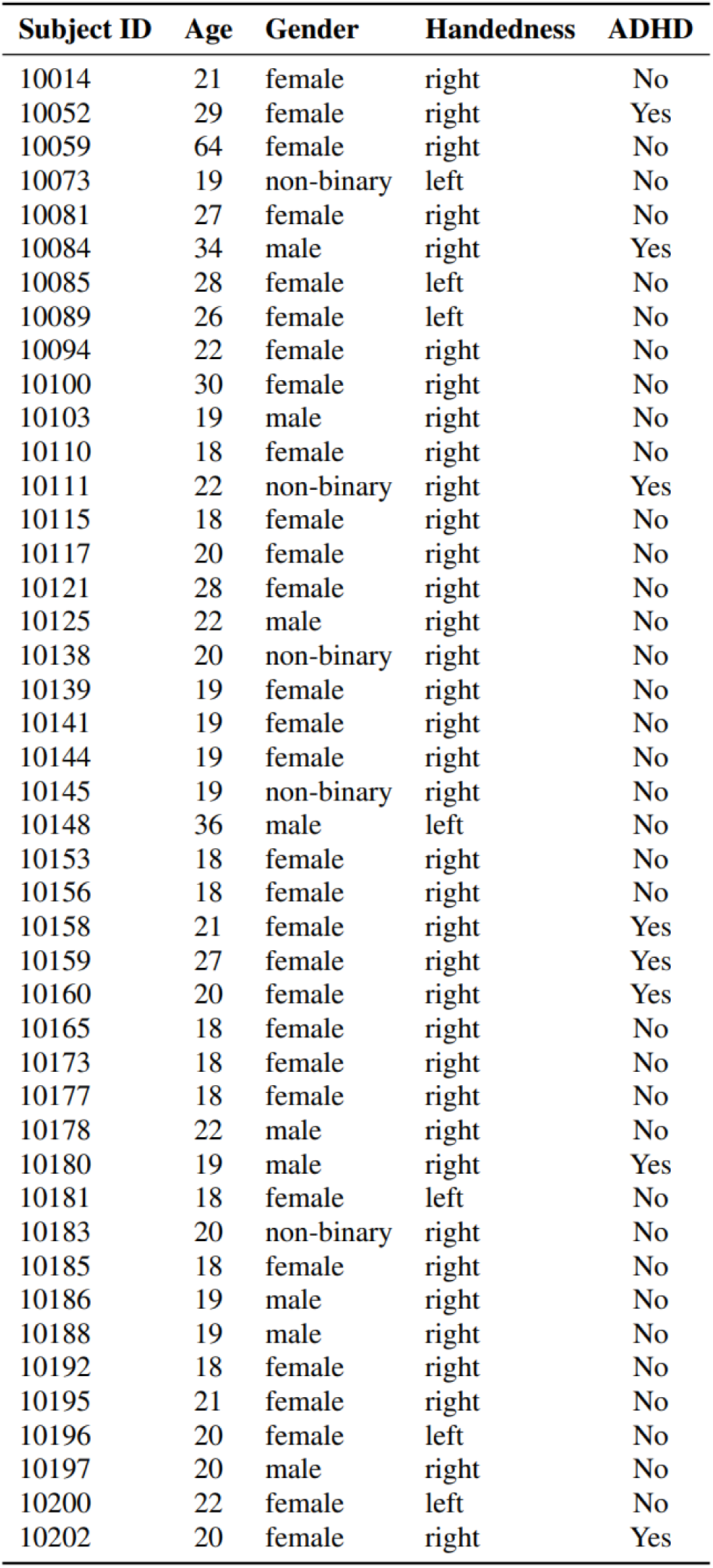
44 participant demographics information.

| **Subject ID** | **Age** | **Gender** | **Handedness** | **ADHD** |
| --- | --- | --- | --- | --- |
| 10014 | 21 | female | right | No |
| 10052 | 29 | female | right | Yes |
| 10059 | 64 | female | right | No |
| 10073 | 19 | non-binary | left | No |
| 10081 | 27 | female | right | No |
| 10084 | 34 | male | right | Yes |
| 10085 | 28 | female | left | No |
| 10089 | 26 | female | left | No |
| 10094 | 22 | female | right | No |
| 10100 | 30 | female | right | No |
| 10103 | 19 | male | right | No |
| 10110 | 18 | female | right | No |
| 10111 | 22 | non-binary | right | Yes |
| 10115 | 18 | female | right | No |
| 10117 | 20 | female | right | No |
| 10121 | 28 | female | right | No |
| 10125 | 22 | male | right | No |
| 10138 | 20 | non-binary | right | No |
| 10139 | 19 | female | right | No |
| 10141 | 19 | female | right | No |
| 10144 | 19 | female | right | No |
| 10145 | 19 | non-binary | right | No |
| 10148 | 36 | male | left | No |
| 10153 | 18 | female | right | No |
| 10156 | 18 | female | right | No |
| 10158 | 21 | female | right | Yes |
| 10159 | 27 | female | right | Yes |
| 10160 | 20 | female | right | Yes |
| 10165 | 18 | female | right | No |
| 10173 | 18 | female | right | No |
| 10177 | 18 | female | right | No |
| 10178 | 22 | male | right | No |
| 10180 | 19 | male | right | Yes |
| 10181 | 18 | female | left | No |
| 10183 | 20 | non-binary | right | No |
| 10185 | 18 | female | right | No |
| 10186 | 19 | male | right | No |
| 10188 | 19 | male | right | No |
| 10192 | 18 | female | right | No |
| 10195 | 21 | female | right | No |
| 10196 | 20 | female | left | No |
| 10197 | 20 | male | right | No |
| 10200 | 22 | female | left | No |
| 10202 | 20 | female | right | Yes |

